# Fast pace of life is associated with host status in small mammals, despite pervasive surveillance bias

**DOI:** 10.64898/2026.06.03.729761

**Authors:** David Simons, Ricardo Rivero, Grant Rickard, Ana Martinez-Checa, Harry Gordon, David W. Redding, Stephanie N. Seifert

**Author notes:** Corresponding Authors: David Simons and Stephanie N. Seifert.

## Abstract

Small mammals are the principal hosts of arenaviruses and hantaviruses, yet the determinants of host status remain contested because global surveillance is non-random. We harmonised 729 studies comprising 695,000 diagnostic assays across 637 species and asked whether host status is predictable once surveillance effort is accounted for.

Surveillance intensity was predicted by night-time light intensity and accessibility, but not local host richness, and sampling effort was phylogenetically clustered (Pagel’s λ = 0.43). Forty-six percent of rodent genera have never been sampled.

In Bayesian phylogenetic mixed models of 43,677 host-virus pairs, faster-lived species tended to be detected as hosts more often, though the credible interval included zero (posterior mean = −0.125, 95% CrI [−0.29, 0.04], pd = 93.2%); the estimate was comparable when restricted to active infection. Synanthropy was independently associated with host status, obligate commensals showing approximately twice the odds. Predictions discriminated hosts from non-hosts in withheld continents (AUC 0.81–0.88), scored at observed sampling effort.

Host and virus phylogenies were congruent in both families, though most associations departed from strict co-divergence, indicating host-switching alongside co-evolutionary history. Host phylogenetic diversity accumulated without saturating, and arenavirus hosts were discovered in a more phylogenetically clustered order than expected by chance, consistent with surveillance concentrating on relatives of known hosts.

Projected globally, community host probability varied largely independently of species richness (R^2^ = 0.028), so diverse assemblages do not carry systematically elevated host probability.

Surveillance bias is substantial and quantifiable, and host biology retains predictive value once it is accounted for.

**Importance:** Arenaviruses and hantaviruses are carried by rodents and other small mammals, outbreaks of infections in humans usually follow contact with these animals. Identifying which species carry them can assist public health agencies in identifying where spillover can occur. However, previous sampling has been very uneven. We show it followed wealth and road access rather than the diversity of small mammals, and that almost half of rodent genera have never been tested for either virus family. Once this unevenness is accounted for, host species remain partly predictable from biology: species that live fast and breed early were somewhat more likely to be recorded as hosts, and species living closely alongside people about twice as likely. Models trained on some continents identified hosts on others. New hosts also tend to be found among close relatives of known hosts, so current host lists should be used as a starting point for future investigations.

## Introduction

Zoonotic spillover is structured by the distribution of reservoir hosts, transmission dynamics within these populations (hazard), opportunities for transmission to susceptible humans (risk), and societal factors that shape susceptibility and response (vulnerability)^1^. Among the most consequential mammalian reservoirs are those hosting viruses in the families *Arenaviridae* and *Hantaviridae*, negative-sense RNA viruses responsible for a spectrum of human diseases including haemorrhagic fevers to severe pulmonary and renal syndromes worldwide^2,3^. While the majority of these zoonoses represent dead-end spillover events, specific lineages (e.g., *Lassa mammarenavirus* and *Andes orthohantavirus*) are capable of sustained human-to-human transmission, elevating them to pathogens of pandemic concern^4–6^. Although these viruses are globally distributed, their public health burden is highly uneven, disproportionately affecting communities with limited healthcare infrastructure, where outbreaks further strain already scarce resources^7–9^. Historically, the understanding of these pathogens focused on a one-host-one-virus paradigm, implying a tight evolutionary co-divergence between virus and host^10–12^. However, recent studies indicate that the ecology and persistence of several arenaviruses and hantaviruses likely involve complex, multi-host networks, in which zoonotic hazard is shaped not just by the presence of a specific host, but by the ecological dynamics of the broader reservoir community^13–16^.

Despite decades of effort, our understanding of reservoir ecology remains distorted. Surveillance is rarely random; it is guided by logistical feasibility, funding, and accessibility creating a surveillance bias analogous to the “streetlight effect”, in which effort concentrates where detection is easiest. Researchers predominantly sample species that are easy to capture (e.g., synanthropes), proximate to research infrastructure (primarily high-income nations), or implicated in known outbreaks as part of reactive public health responses^17–19^. This results in a surveillance landscape that reflects human activity more than viral distribution. This distortion need not be limited to imprecision in the ecological signal. If the same socioeconomic drivers of surveillance effort (e.g., wealth, accessibility, night-light intensity) are independently correlated with host ecological traits, the resulting non-random sample can function analogously to conditioning on a collider variable, inducing or masking an association between host traits and detected host status that need not reflect the underlying biological relationship^20^. Synanthropic, fast-lived species, for example, may be disproportionately represented in accessible, high-income regions for reasons unrelated to any causal role in viral maintenance, such that their apparent association with host status could be inflated, diminished, or reversed relative to a representative sample. As a result, the apparent association between host status and commensal rodents (e.g., *Rattus* spp., *Mus* spp.) may reflect detection bias as much as biological reality^21,22^. Here, we argue that disentangling the ecology of these viruses is impossible without first quantifying and statistically adjusting for this anthropogenic filter.

Once surveillance bias is accounted for, macroecological and evolutionary theory offer competing hypotheses for what determines host status. Life-history theory predicts trade-offs between reproductive investment and somatic maintenance: species that mature early and produce large litters may invest less in energetically costly adaptive immunity, favouring pathogen tolerance over resistance^23,24^. This tolerance phenotype may limit disease severity despite high pathogen burdens, compromising viral clearance and promoting persistent infection that facilitates transmission^25^. The ecological opportunity hypothesis instead proposes that generalists with broad diets, large ranges and wide habitat tolerances encounter more diverse pathogens, increasing rates of acquisition and of novel host-virus associations^26^.

At the community level, these traits shape landscape-level hazard. We use “ecological filtering” for the combined abiotic and biotic processes shaping host-assemblage composition, broader than “environmental filtering”, which is restricted to abiotic tolerance^27^. Whether this filtering consistently favours high host-probability generalists in species-poor, human-dominated environments, elevating local host probability relative to intact ecosystems, remains unresolved^28,29^. Evolutionary history also matters: strict co-speciation implies restricted host ranges constrained by deep time, while phylogenetic incongruence would indicate host-switching, facilitated by ecological contact, as a driver of viral diversity^30^. Recent frameworks have called for integrating such macroecological properties into zoonotic risk modelling^31^.

Identifying reservoirs from historical surveillance is inherently fraught. The detection of a pathogen (through seroconversion, molecular detection, or viral isolation), does not definitively resolve a species’ role in chains of transmission. An animal may be a true maintenance reservoir, an incidental dead-end host, or merely share an environment with the true reservoir. Rather than attempting to assign rigid, binary reservoir classifications to historically messy data, we use the term “host” to refer to any species demonstrating evidence of viral infection (via PCR, sequencing, or isolation) or seroconversion (detectable antibodies reactive to the target virus, noting that polyclonal cross-reactivity means such assays support specificity at approximately viral family level rather than to a single virus species), without implying a specific epidemiological role such as reservoir or maintenance status. We posit that across 50 years of global data, consistent patterns of seroconversion and infection will reveal the underlying ecological networks of these viruses, provided that severe anthropogenic sampling biases are mathematically controlled.

Responding directly to this gap, this study aims to move beyond descriptive surveillance mapping to a mechanistic understanding of reservoir ecology. By synthesising over 50 years of global surveillance data, we first quantify the anthropogenic filter, mapping the taxonomic, geographic, and temporal biases in small mammal sampling. We then employ Bayesian phylogenetic models to test whether pace of life and synanthropy predict host status after adjusting for sampling effort. Concurrently, we assess the evolutionary congruence between hosts and viruses to determine the relative roles of host-switching versus co-divergence. Finally, we leverage these models to project the host potential of unsampled species, identifying potential hotspots of hazard that current surveillance has missed. This approach allows us to ask where hosts are likely to occur, independent of where they have been looked for.

## Results

### The Anthropogenic Filter: Quantifying Surveillance Bias

Global surveillance for *Arenaviridae* and *Hantaviridae* in small mammals is fundamentally decoupled from host biodiversity, driven instead by socioeconomic and geographic convenience. Our systematic review extracted field records for over 577,000 observed small mammals. Of these, approximately 561,000 individuals were subjected to virological screening, yielding nearly 695,000 distinct assay results across 637 host species. Despite this volume of data, the distribution of these records reveals a highly uneven surveillance effort. After 50 years of research, 46% of all Rodentia and Eulipotyphla genera remain unsampled, with entire families such as *Gliridae* receiving negligible surveillance attention (Supplementary Figure S2). In our Zero-Inflated Negative Binomial (ZINB) model of global surveillance (Fig. 1; Supplementary Table S1), the strongest predictors of sampling intensity at the district level were night-light intensity and accessibility. Since accessibility was defined as travel time to major cities, this negative coefficient indicates that surveillance intensity substantially decreases as locations become more remote (Supplementary Figure S3). Conversely, neither local species richness nor human population density were important predictors of sampling effort. These coefficients describe sampling intensity where surveillance occurred. In the zero-inflation component, which models whether a district was sampled at all, higher species richness was associated with a lower probability of a district being unsampled, and higher population density with a greater probability (Supplementary Table S1), although these estimates are less precisely identified than those of the count component.

**Fig. 1.**
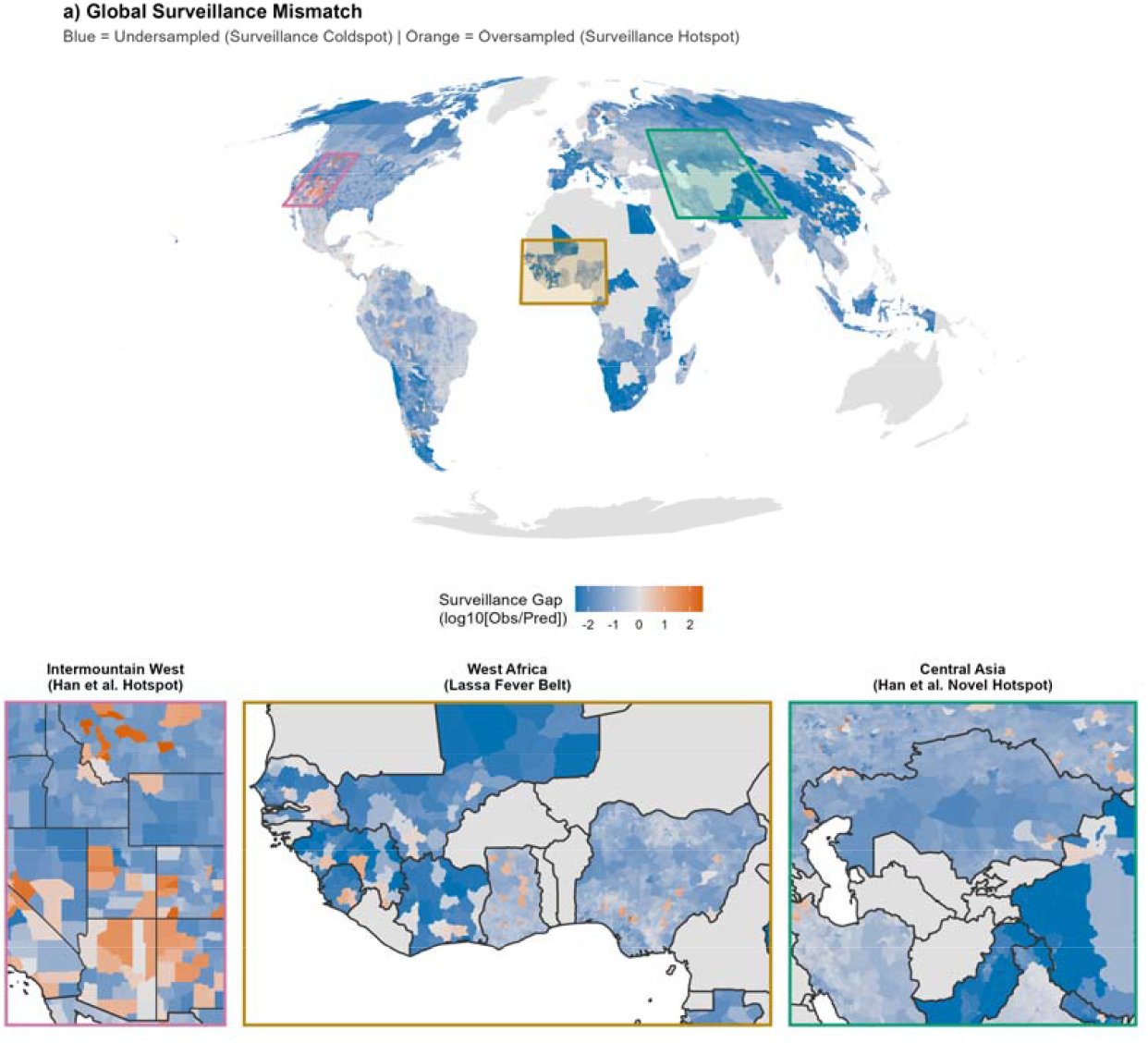
Global surveillance inequality. Global map of surveillance residuals from a Zero-Inflated Negative Binomial (ZINB) model. Colours indicate the log-ratio of observed versus predicted surveillance effort (log10 [Observed/Predicted]). “Hotspots” (Orange, >0) represent districts sampled more intensely than predicted by night-light intensity, accessibility, population density, and local host richness; “Coldspots” (Blue, <0) represent undersampled regions. Insets highlight key regional anomalies: the Intermountain West (oversampled), West Africa (targeted Lassa surveillance), and Central Asia (undersampled).

These geographic constraints generate taxonomic distortions. Surveillance is heavily skewed towards the Palearctic and Nearctic realms, which collectively account for 71% of all detected small mammals, leaving a substantial number of the most biodiverse regions in the tropics largely uncharacterised. At the genus level, sampling coverage is driven primarily by geographic ubiquity rather than evolutionary distinctiveness (Supplementary Figure S4). We observed a strong positive relationship between total geographic range size and the area sampled; genera with massive distributions (e.g., *Rattus, Mus, Apodemus*) are sampled across broad spatial extents, whereas range-restricted genera are frequently ignored. However, even for widespread genera, the proportional coverage remains low; despite *Mus* being sampled in 444 administrative districts, this represents less than 3.3% of its IUCN recorded range.

Generalised Additive Models (GAMs) showed that host body mass and geographic range size were the dominant predictors of species-level sampling effort (*p* < 0.001), with larger and more widespread species receiving disproportionate attention. While totally synanthropic species exhibited higher mean sampling estimates than non-synanthropic species, this effect did not reach statistical significance likely due to the rarity of this trait in the global dataset (*N* = 6 of 2,436 modelled species). Instead, geographic range size remained the single strongest driver of surveillance effort (*χ*^2^ = 422.7). Sampling effort also showed significant phylogenetic signal (Pagel’s λ = 0.43, likelihood-ratio test vs. λ = 0: *p* < 0.001), indicating that closely related host species tend to receive similar levels of surveillance attention rather than sampling being phylogenetically random.

Temporally, surveillance is fundamentally reactive. Global sampling effort exhibits distinct pulses following major zoonotic discovery events, most notably the description of Lassa virus (1969) and the Sin Nombre virus outbreak (1993). GAMs reveal that while surveillance in the Americas grew exponentially since the mid-1990s (*p* < 0.001) before declining post 2005, effort in Africa and Asia has remained comparatively static and highly episodic, driven by short-term outbreak responses rather than sustained monitoring.

### Macroecological Associations with Host Status

We disentangled intrinsic biological suitability from the surveillance biases identified above by fitting Bayesian phylogenetic generalised linear mixed models (GLMMs) to the dyadic host-virus data (*N* = 43,677 dyadic pairs). As expected, sampling effort was the dominant predictor of host status; the number of individuals tested had a strong positive effect on the probability of detection (Supplementary Table S2), indicating that widely detected hosts are often the most intensely scrutinised species (Fig. 2a). After adjusting for sampling effort and phylogeny, the first principal component of life history (PC1), representing the slow-fast continuum, showed a negative association with host status (posterior mean = −0.125, 95% CrI [−0.29, 0.04], pd = 0.932). Lower PC1 scores correspond to smaller body mass, earlier maturity and larger litter sizes, so this indicates fast-lived species were more likely to be detected as hosts. The credible interval includes zero. Restricting positive dyads to active infection only (PCR, sequencing, or viral isolation) produced an estimate of similar magnitude (posterior mean = −0.17 [−0.40, 0.07], pd = 91.8%), consistent with the association not being an artefact of serological cross-reactivity alone.

**Fig. 2.**
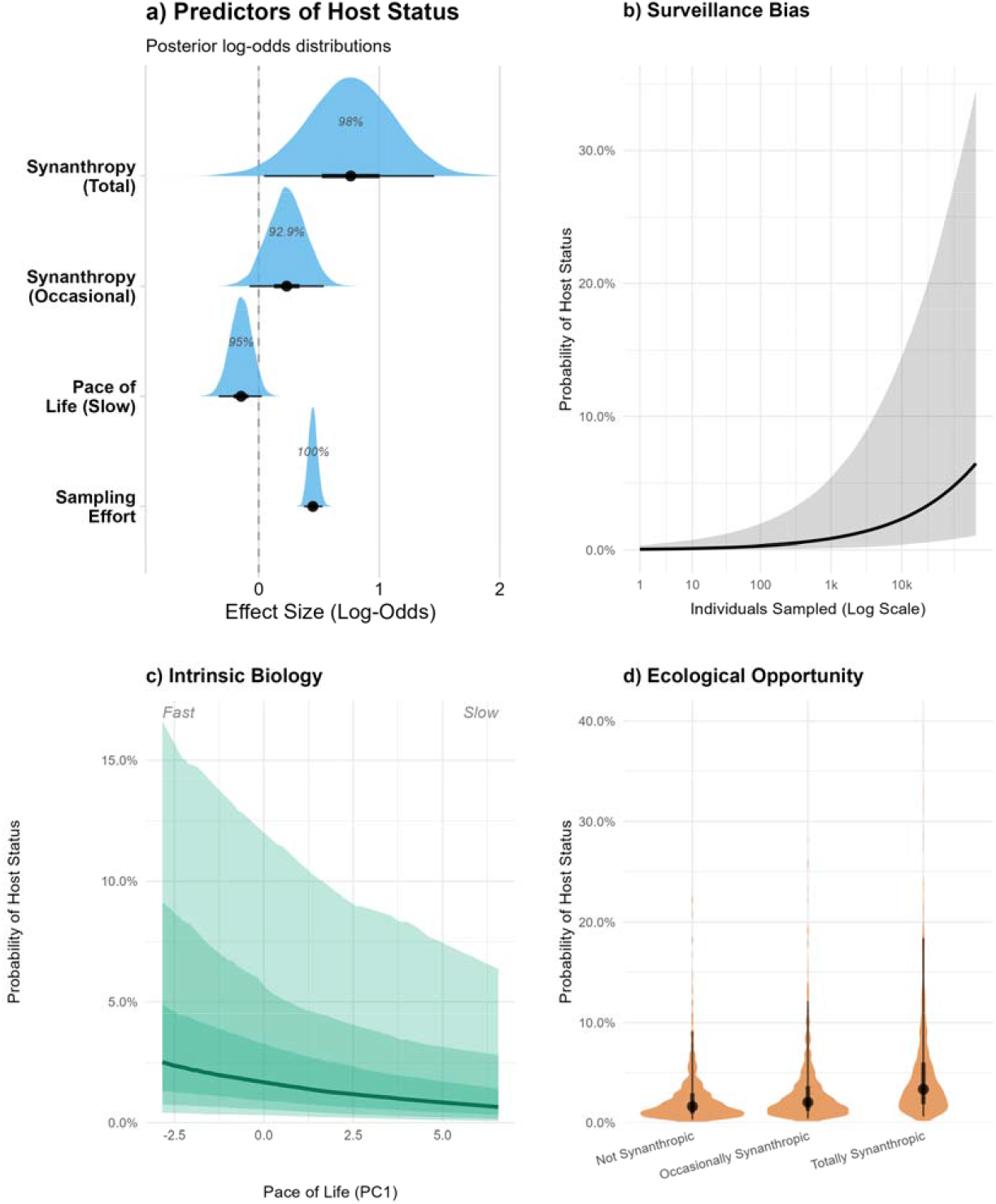
Associations with host status. a) Posterior log-odds distributions for the factors associated with host status in a Bayesian phylogenetic GLMM (*N* = 18,906). Percentages indicate the Probability of Direction (pd), representing the certainty that the effect is non-zero in the indicated direction. Note that these are derived from the synanthropy subset model and therefore differ from values reported for the global model in the main text. b)–d) Marginal effects of the predictors on the probability of being a host. (b) Sampling effort is the strongest driver of discovery; the curve visualises probability across the observed range of sampling effort, shaded areas represent 95% Credible Intervals. c)–d) Intrinsic biological drivers visualised conditional on high (95th percentile) observed sampling effort. c) Fast life history (low PC1 scores) is associated with a higher probability of host status, shaded bands represent the 50%, 80% and 95% credible intervals. d) Totally synanthropic species (obligate commensals) exhibit higher host probability than non-synanthropic species, highlighting the role of ecological opportunity, intervals represent 50% and 95% credible intervals.

In a subset analysis accounting for human affinity (*N* = 18,906 dyadic pairs), synanthropy was a distinct predictor of host status independent of life history (Fig. 2a). Totally synanthropic species (obligate commensals) had a higher probability of detected host status than non-synanthropic species (posterior mean = 0.762, 95% CrI [0.045, 1.46]; Supplementary Table S3), an approximate 2.2-fold increase in odds, though this interval only narrowly excludes zero. Occasionally synanthropic species showed a weaker effect. The pace-of-life coefficient was essentially unchanged by the addition of the synanthropy term (posterior mean −0.147 in this subset versus −0.125 in the global model), indicating the two traits contribute largely non-redundant information rather than re-detecting a single underlying signal (Fig. 2c and Supplementary Figure 12)

Partitioning the model variance (Supplementary Table S2) indicated that host status is a labile trait. The random effect variance attributed to specific host identity was four times larger than the variance attributed to phylogenetic history, indicating that host status is highly species-specific rather than a deeply conserved phylogenetic trait. Furthermore, we observed substantial variation in detection probability among viral species. Post-hoc interrogation of the viral random effects revealed no strong evidence of a systematic difference in baseline detectability between the two viral families (posterior mean difference in family-level random intercepts = −0.27 [95% CI: −0.58, 0.03]; Supplementary Figure S5).

We conducted a sensitivity analysis to confirm these patterns were not artefacts of historical serological cross-reactivity or dead-end spillover events, restricting positive host status exclusively to molecular detection or viral isolation. In this strict model, the pace-of-life estimate was comparable to the full dataset containing serology (Supplementary Table S4 vs. S2). The credible interval widened owing to the reduction in positive detection events, but the direction and magnitude were consistent, indicating the association is not driven by exposure proxies alone. Restricting the data to molecular evidence reduced the variance attributed to specific viral species while increasing idiosyncratic host-level variance.

A subsequent complete-case analysis ensured the observed association between a fast pace of life and host status was not an artefact of phylogenetic imputation. Refitting the model to a restricted dataset containing only empirically recorded trait values (*N* = 14,697 dyadic pairs) yielded a stable posterior mean (Supplementary Table S5). Variance partitioning within this empirical subset supported that species-specific variance outweighed phylogenetic variance in the absence of imputed covariance. Together, these parallel models (Supplementary Tables S2-S5; Supplementary Figure S6) show that the direction and approximate magnitude of the pace-of-life association are consistent across evidentiary definitions and trait-imputation approaches, indicating the signal is not an artefact of historical serological noise or assumed phylogenetic proximity.

Scaling these individual traits to the landscape level, we projected these estimates spatially to map the global distribution of community host probability (Fig. 3). We first executed a Leave-One-Region-Out (LORO) spatial cross-validation, iteratively withholding each continent and predicting host status for the withheld species using only the biological relationships learned elsewhere. Predictions discriminated hosts from non-hosts in withheld continents with an area under the receiver operating characteristic curve (AUC) of 0.810 (Americas), 0.832 (Asia), 0.826 (Europe), and 0.876 (Africa), scored against observed host status at each dyad’s actual sampling effort. Genus-level overlap between withheld and training sets was low in most folds (8.8% Americas, 13.0% Africa, 26.7% Asia), though higher for Europe (37.5%), the smallest fold, where Old World genera span multiple continents. Coefficient estimates were also stable across folds: out-of-sample predictions correlated with full-model estimates at Spearman’s *ρ* = 0.767 (*p* < 0.001). Regional comparison of these out-of-sample predictions indicated that East Asian assemblages had the highest median predicted host probability (0.0434), higher than the Atlantic Forest, North America, the unclassified global baseline, and West Africa (all pairwise *p* <5 × 10^−11^; Supplementary Figure S7). The remaining regions were largely indistinguishable. Western Europe was excluded from this comparison, as only three species met the majority-occurrence criterion for regional assignment. We present these comparisons as a check on out-of-sample model behaviour rather than as regional estimates of host probability.

**Fig. 3.**
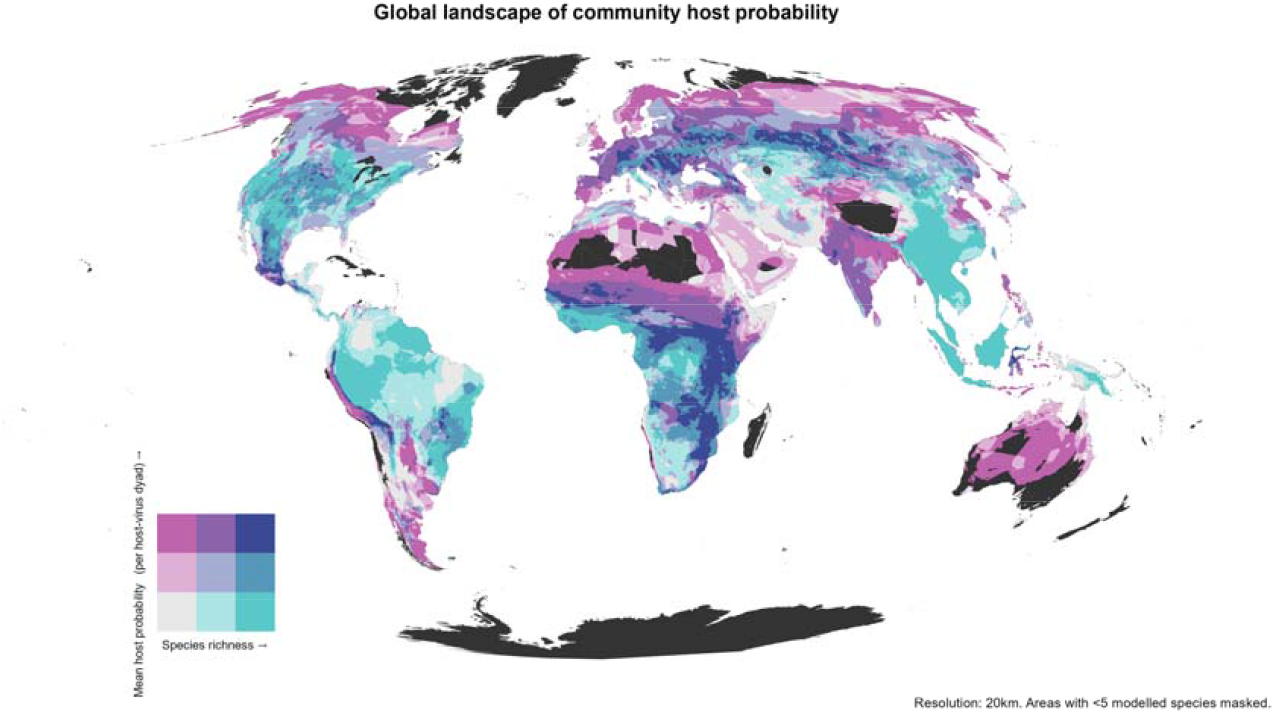
The global landscape of community host probability. Bivariate map of species richness (x-axis) and community host probability (y-axis) at 20km resolution. Community host probability is the mean predicted probability of host status across small mammal species present in each cell, derived from the global Bayesian GLMM using pace-of-life traits (*N* = 2,766 species); values are per host-virus dyad, evaluated at the 95th percentile of observed sampling effort, not the probability that a species is a host of arenaviruses or hantaviruses generally. The two axes are near-independent across districts (*R*^2^ = 0.028), the classification therefore represents two separate components of community-level hazard. High values of mean host probability occur predominantly in species-poor arid and montane systems (e.g., the Sahel, Andean highlands): averaging across few species can raise the mean where assemblages are small. Districts combining high richness with high host probability (upper right) include Cochabamba (Bolivia), Oaxaca (México) and Turkana (Kenya). Cells with fewer than 5 modelled species are masked. Synanthropy is not included in this projection, as data are unavailable for most non-target species.

To maximise taxonomic coverage for the final map, we utilised the global in-sample model (*N* = 43,677) to predict host probability for all small mammal species for which life history data were available (*N* = 2,766).

Bivariate classification indicated that the highest mean community host probability occurred in species-poor rather than species-rich assemblages (Class 1-3: mean host probability 0.0392, mean richness 12.1 species; Class 3-1: 0.0348, 29.4 species), a 13% relative difference between the extremes. A subset of districts combined high richness with high host probability (Class 3-3: 0.0387, 26.4 species). Across all districts, however, species richness explained little variation in mean host probability (linear model slope = −4.5 x 10 ^− 5^ per species, *R*^2^ = 0.028, *N* = 37,676).

Representative districts for each class are given in Table 1. The complete dataset of predicted community host probability and species richness for 37,676 administrative districts is available as a supplementary data file.

**Table 1:**
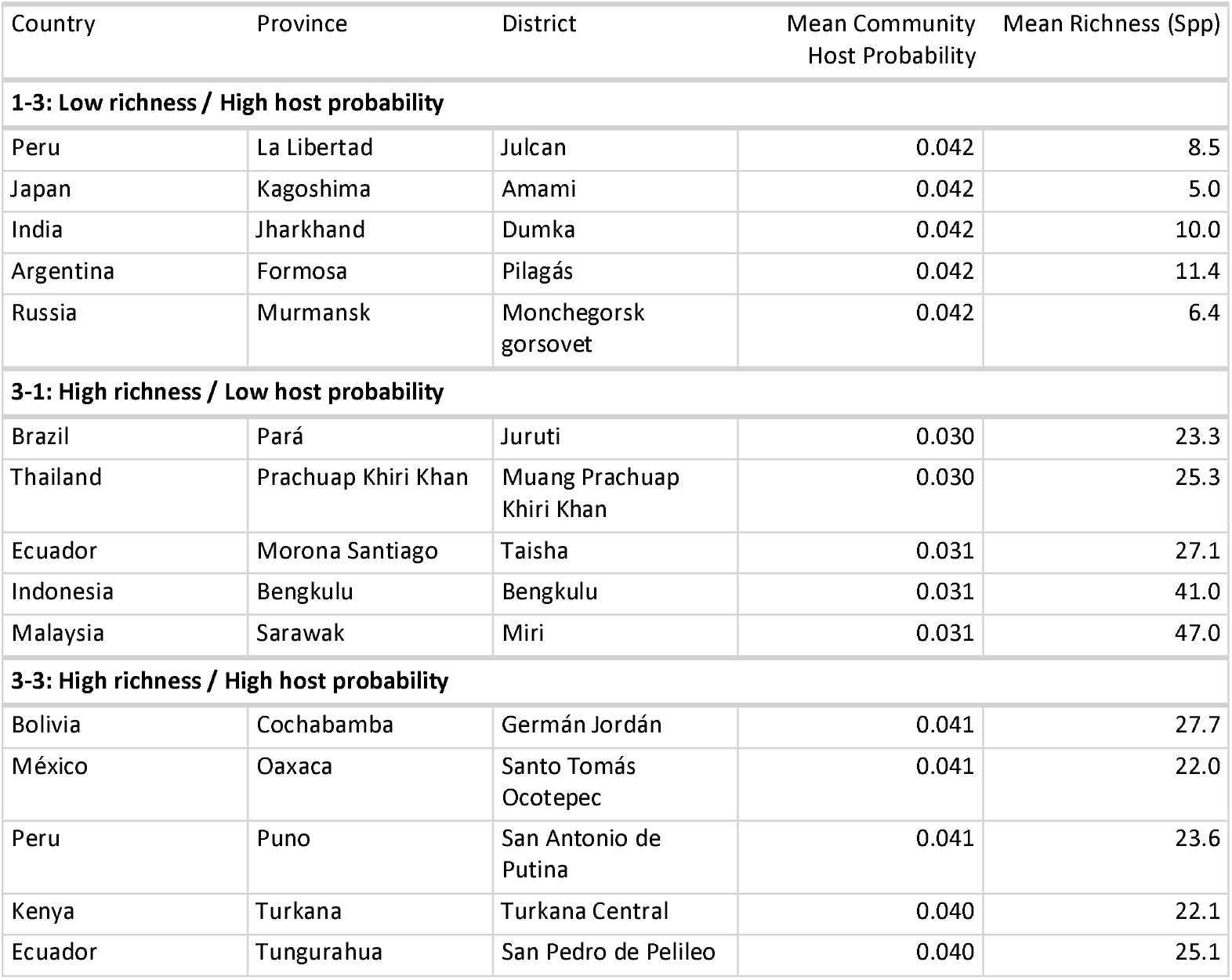
Representative districts for the three bivariate risk classes.

| Country | Province | District | Mean Community Host Probability | Mean Richness (Spp) |
| --- | --- | --- | --- | --- |
| **1-3: Low richness / High host probability** |  |  |  |  |
| Peru | La Libertad | Julcan | 0.042 | 8.5 |
| Japan | Kagoshima | Amami | 0.042 | 5.0 |
| India | Jharkhand | Dumka | 0.042 | 10.0 |
| Argentina | Formosa | Pilagás | 0.042 | 11.4 |
| Russia | Murmansk | Monchegorsk gorsovet | 0.042 | 6.4 |
| **3-1: High richness / Low host probability** |  |  |  |  |
| Brazil | Pará | Juruti | 0.030 | 23.3 |
| Thailand | Prachuap Khiri Khan | Muang Prachuap Khiri Khan | 0.030 | 25.3 |
| Ecuador | Morona Santiago | Taisha | 0.031 | 27.1 |
| Indonesia | Bengkulu | Bengkulu | 0.031 | 41.0 |
| Malaysia | Sarawak | Miri | 0.031 | 47.0 |
| **3-3: High richness / High host probability** |  |  |  |  |
| Bolivia | Cochabamba | Germán Jordán | 0.041 | 27.7 |
| México | Oaxaca | Santo Tomás Ocotepec | 0.041 | 22.0 |
| Peru | Puno | San Antonio de Putina | 0.041 | 23.6 |
| Kenya | Turkana | Turkana Central | 0.040 | 22.1 |
| Ecuador | Tungurahua | San Pedro de Pelileo | 0.040 | 25.1 |

### Evolutionary Congruence and Host-Switching

While ecological filtering shapes the spatial distribution of these host communities, the assembly of these multi-host networks is driven by a complex evolutionary history. Host-virus associations both in *Arenaviridae* and *Hantaviridae* are characterised by broad lineage-level co-divergence punctuated by frequent, recurrent host switching. By comparing host and viral phylogenies with family-specific incidence matrices, we found significant cophylogenetic structure in both families. For *Arenaviridae* (16 viruses, 41 hosts, 47 links), the Procrustean Approach to Cophylogeny (PACo) detected significant congruence (*p* < 0.001, 999 permutations) as did ParaFit (*p* = 0.001). For *Hantaviridae* (30 viruses, 90 hosts, 141 links), PACo also showed significant global congruence (*p* < 0.001, 999 permutations), consistent with the ParaFit global result (*p* = 0.001).

Concurrently, link-level PACo residuals were broadly dispersed in both families (*Arenaviridae* median residual = 31.8; *Hantaviridae* = 45.0), with clusters of high-congruence links coexisting with diffuse low-congruence links (Fig. 4), supporting a model in which broad lineage-level structure co-occurs with recurrent host-switching at finer taxonomic scales. These results indicate that while host-virus associations retain significant lineage-level evolutionary structure, frequent link-level incongruence reveals recurrent cross-species transmission beyond strict co-divergence. This mixed evolutionary pattern suggests that observed host ranges are not rigidly fixed by deep evolutionary time, motivating a temporal assessment of realised host breadth.

**Fig. 4.**
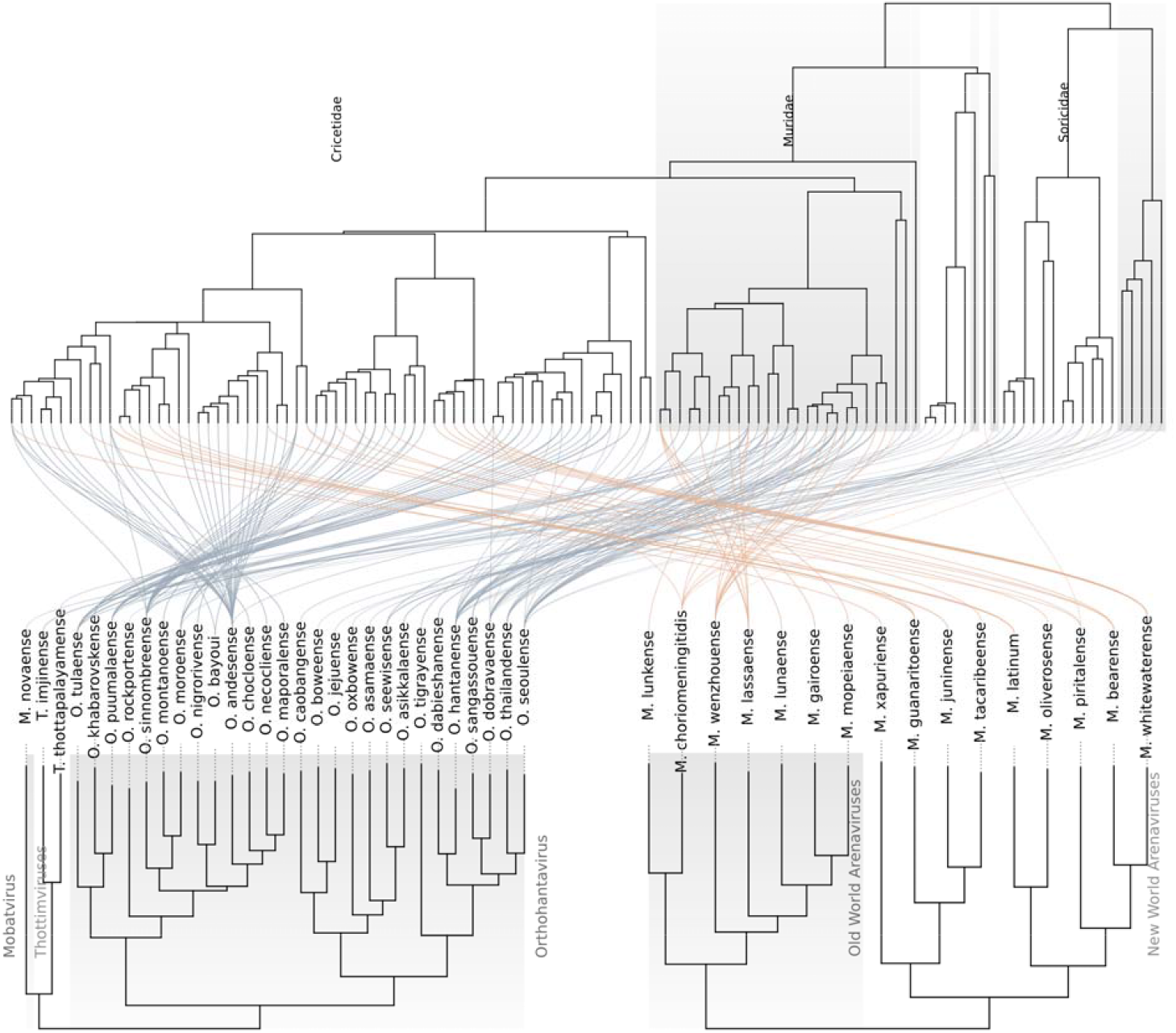
Phylogenetic congruence between *Hantaviridae* and *Arenaviridae* and their mammalian hosts. Tanglegram mapping the mammalian host phylogeny (top) against viral phylogenies for *Hantaviridae* (bottom left) and *Arenaviridae* (bottom right). Curved links connect each virus to PCR-positive host species; link width and opacity are scaled inversely with PACo residuals, thicker, darker links indicate stronger phylogenetic congruence and thinner, fainter links indicate weaker congruence (host-switching). Host background bands denote mammalian families, and viral bands denote viral genera (*Mobatvirus, Thottimvirus, Orthohantavirus*) or major clades (Old World and New World arenaviruses). Hantavirus links are concentrated on Cricetidae, aside from shrew-borne *Mobatvirus* and *Thottimvirus*; arenavirus links largely separate Old World viruses on Muridae from New World viruses on Cricetidae.

### Host Discovery Is Phylogenetically Clustered and Has Not Saturated

To quantify how the mixed evolutionary mosaic of co-divergence and host-switching translates into realised host breadth over time, we analysed the cumulative phylogenetic diversity (*PD*_*c*_) of infected hosts at the family and viral species levels. The maximum annual gain in host phylogenetic diversity (*ΔPD*_t_) increased through time in *Arenaviridae* (Spearman *r* _*max*_= 0.39, one-tailed *p* = 0.029) but not in *Hantaviridae* (*r* _*max*_ = 0.01, one-tailed *p* = 0.49). Across all virus-year observations, correlations were weakly positive in both families (*Arenaviridae*: *r*_*all*_ = 0.27; *Hantaviridae*: *r* _*max*_ = 0.18), indicating that newly detected hosts continued to add phylogenetic diversity.

Family-pooled cumulative curves also showed no saturation (Fig. 5c and d). After pooling hosts by family and ordering them by first detection year, *Arenaviridae* reached 44 hosts with a *PD*_*c*_ Family-pooled cumulative curves also showed no saturation (Fig. 5c and d). After pooling hosts of 919.5 and *Hantaviridae* reached 95 hosts with a *PD*_*c*_ of 1,593.4. These values correspond to 46.8% and 81.0% of the total branch length of the analysed host tree. Compared with 10,000 random orderings of the same hosts, arenavirus hosts were detected in a more phylogenetically clustered sequence than expected (area under the observed curve 13% below the null median, one-tailed *p* = 0.037). Close relatives of known hosts were found first, followed by more divergent lineages. Hantavirus discovery showed a similar tendency but did not reach significance (4% below the null median, *p* = 0.12).

**Fig. 5.**
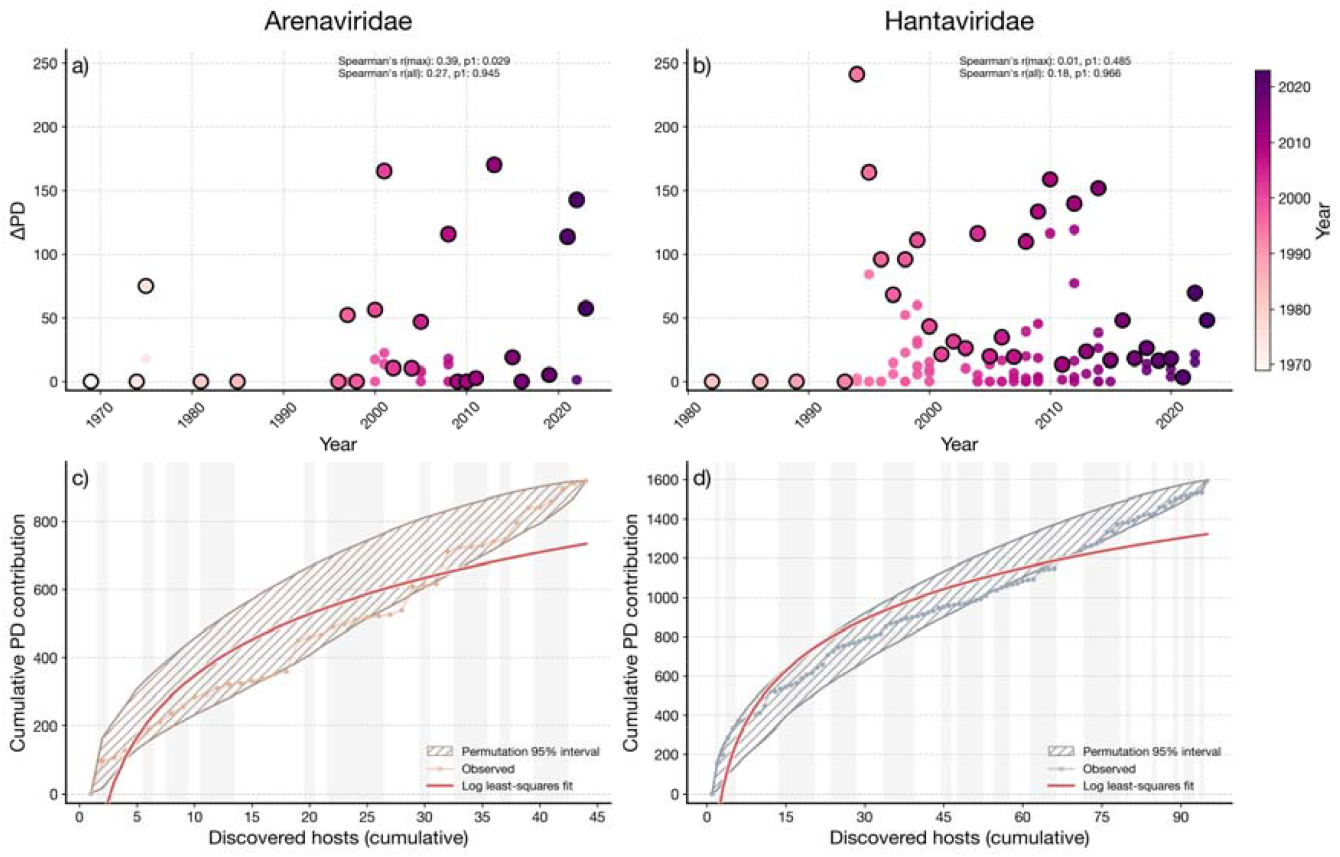
Family-level accumulation of host phylogenetic diversity. a) Annual change in host phylogenetic diversity () for arenaviruses. Each point is a host-year discovery event, coloured by calendar year; outlined circles indicate the yearly maximum event. b) Annual for hantaviruses, with the same point encoding and yearly maximum outlines. Spearman trend statistics for yearly maxima () and all points () are shown in a) and b). Both *p*-values are one-tailed, tested for an increase through time and tested for a decrease, so that a high *p*-value for indicates no evidence that newly detected hosts became phylogenetically redundant. c) Cumulative host for arenaviruses as hosts are added in empirical discovery order. The hatched band is the 95% null envelope from 10,000 random permutations of host discovery order; the observed trajectory is shown as the coloured line and points, and the red curve is the logarithmic least-squares fit. d) Cumulative host for hantaviruses, shown as in c). Light grey vertical bands in c) and d) mark successive discovery-year intervals. Where the observed path lies along or below the lower bound, hosts were added in a more phylogenetically clustered order than expected from a random discovery sequence.

Across individual viral species, we detected strong heterogeneity in host breadth. We classified each virus into one of three trajectory archetypes using rank-based combinations of host richness, *PD* per host, and relative jump magnitude. We classified 30 viruses as having sparse or sampling-limited histories (B-type), 18 as showing punctuated bursts (C-type), and one as a sustained accumulator (A-type). All 30 B-type viruses had ≤2 hosts and ≤2 year batches. For many of these viruses, apparently narrow host ranges may therefore reflect limited sampling rather than fixed biological limits.

We also examined six exemplar viral trajectories (Supplementary Figure S8), whose records spanned 7 to 21 hosts and 4 to 12 year batches, to assess this family-level pattern in individual viruses. For some viruses, the largest jumps followed recognised human outbreaks, as with Sin Nombre, for which six new hosts were detected in 1994. For others, the jumps were more consistent with rodent-focused field campaigns (see Supplementary Text for detailed epidemiological contextualisation of these trajectories).

## Discussion

Our synthesis indicates that the prevailing understanding of *Arenaviridae* and *Hantaviridae* reservoir ecology is distorted by an anthropogenic filter. Global surveillance effort is driven not by host biodiversity or ecological relevance, but by logistical convenience and socioeconomic wealth: night-light intensity and accessibility predict where sampling occurs, while local host richness has no credible effect. Despite this distortion, host status retains predictable structure once sampling effort is accounted for. Our models discriminated hosts from non-hosts in withheld continents with an AUC of 0.81–0.88, scored at each dyad’s observed sampling effort, indicating that the fitted relationships transfer beyond the regions where they were learned. Because effort contributes to these predictions, and because genus-level overlap between withheld and training sets was not zero, this should not be read as discrimination from biology alone. Within that signal, a fast pace of life emerges as a modest, individually uncertain, but directionally consistent correlate of host status, alongside a stronger association with synanthropy. Our community-level projections are consistent with anthropogenic landscapes supporting assemblages shifted toward fast-lived generalists with higher host probability^21^, indicating that pristine tropical ecosystems do not carry systematically higher host probability than species-poor assemblages.

This pervasive “streetlight effect” reflects a legacy of wealth-biased research investment^32^. Because infrastructure and funding remain concentrated in the Palearctic and Nearctic realms, speciose tropical regions remain largely uncharacterised^33^, creating a self-reinforcing taxonomic blind spot in which familiar, accessible lineages are targeted while entire families go unexplored^34,35^. This uneven foundation is further destabilised by a persistent genetic data gap. Over 67% of historical host-virus associations rely solely on serological or fragment-based evidence, which risks conflating viral maintenance with environmental exposure^36,37^. Without lineage-specific sequencing, apparent host ranges are likely artificially inflated. Geographic discontinuities in sampling coverage may generate similar artefacts in viral phylogeography, where genetically continuous populations appear as distinct lineages^38^. Further, surveillance is often targeted toward one viral family over the other by region (e.g., 89.7% *Hantaviridae* in the United States vs. 100% *Arenaviridae* in Nigeria and Sierra Leone), so species-level testing volume may not capture family-specific detection opportunity, a confound that plausibly contributes to the near-null family-level detectability difference reported above.

Establishing a definitive epidemiological threshold separating an incidental host from a true maintenance reservoir remains a persistent challenge in disease ecology^39,40^. Because historical surveillance lacks the longitudinal prevalence data required to prove onward transmission, our framework eschews arbitrary classification cut-offs. Instead, by modelling a macroecological continuum of host status, we identify traits associated with detected infection and exposure; whether these traits also confer viral tolerance, as hypothesised below, remains to be directly tested.

The accumulation of host phylogenetic diversity over time shows that global surveillance is phylogenetically concentrated rather than distributed evenly across candidate hosts. Host breadth has not saturated in either family, so current host lists are best treated as lower bounds. Arenavirus hosts entered the record in a phylogenetically clustered order, with close relatives of known hosts detected before more divergent lineages, consistent with surveillance near known hosts. For some viruses, bursts of host discovery aligned with major public health crises, suggesting that sampling effort is at times mobilised in response to human spillover events^41,42^. Because human encroachment simultaneously drives initial ecological contact, spillover risk, and the healthcare infrastructure required to detect an outbreak, post-outbreak surveillance disproportionately targets anthropogenic landscapes. This contextualises the evolutionary mosaic observed in the cophylogenetic analyses. While *Arenaviridae* and *Hantaviridae* exhibit broad lineage-level co-divergence with their mammalian hosts, link-level residuals reveal frequent, recurrent host-switching^13,43^. Because surveillance typically intensifies only post-outbreak, the available data disproportionately capture the novel host-virus associations that arise when viruses jump into the adaptable, synanthropic host networks already thriving at these human interfaces.

Concurrently, the fast pace-of-life signal is consistent with macroecological hypotheses regarding immune trade-offs, under which fast-lived species are proposed to invest fewer resources in costly adaptive immunity^24^. The directional evidence for this association was comparable under our strictest evidentiary definition, restricted to active infection (pd = 91.8%, versus 93.2% in the primary model), indicating the pattern is not solely attributable to serological cross-reactivity. Whether this reflects reduced immune investment favouring viral tolerance over clearance, and thus the persistent, subclinical infections proposed to support host maintenance^44^, cannot be determined from these surveillance data, and would require longitudinal or experimental infection studies. The high species-specific variance in the models indicates host status is a labile trait, emerging idiosyncratically rather than uniformly across deep evolutionary time. This has been attributed to receptor compatibility^45^, though receptor structure would itself be expected to carry phylogenetic signal; the low phylogenetic variance we observe therefore points to local ecological exposure as the more probable driver.

Several non-exclusive mechanisms likely underlie the observed synanthropy effect. Part of this may reflect an availability bias: commensal-associated viruses, being closer to humans, have had greater historical opportunity to be identified as zoonotic^46^. This is unlikely to fully explain the pattern since the synanthropy effect’s credible interval remained clear of zero, albeit narrowly, after controlling for cumulative sampling effort (Table S3). A subtler discovery-order effect where commensal pathogens are recognised as zoonotic earlier due to their proximity to reactive, outbreak-driven surveillance cannot be excluded given the sparsity of our data in space and time. Beyond sampling artefacts, the ecological correlates of commensalism (e.g., high population density and close spatial association with humans) plausibly contribute to this pattern^22^, though evaluating their relative role in transmission dynamics is beyond the scope of the present species-level analysis.

Scaling these individual traits to the community level, species richness and mean community host probability varied largely independently across global districts. Highly diverse tropical forests, often assumed to be primary frontiers of pandemic risk, showed slightly lower mean host probability (0.0348) than species-poor assemblages (0.0392), though richness explained little of the overall variation. High biodiversity therefore does not itself confer elevated community host probability; neither does it systematically buffer it. Anthropogenic landscapes may support assemblages shifted toward fast-lived species^19,29^, though land-use change was not measured here.

We suggest that districts combining high richness with high host probability (Class 3-3) require particular attention. The cophylogenetic analysis indicates host-switching contributes substantially to the current host-virus network alongside co-divergence; assemblages containing many host species offer more opportunities for such switching, and for the emergence of novel associations. We therefore present it as a hypothesis derived from our projections. By area, these districts concentrate in East and Southern Africa (Uganda, 92% of assessed area; Tanzania, 69%; Zambia, 64%) and in Central and Eastern Europe (Slovakia, 72%; Romania, 66%), alongside the Central Asian steppe. The latter corresponds to a region previously identified as a concentration of rodent reservoir species on the basis of life-history traits^47^. The Intermountain West, also identified in that work, is largely absent from this class in our projections.

Empirical trait data are sparse across tropical realms: mean imputation across our seven life-history traits was 58.5% for Afrotropic, 69.5% for Neotropic, 71.8% for Indomalayan and 73.7% for Australasia, against 32.2% for Nearctic and 6.1% for Palearctic-North, a gap sometimes termed the “Eltonian shortfall”^48^. Predictions for the African and Neotropical districts identified above therefore rest more heavily on phylogenetic imputation than those for temperate regions. Because imputation shrinks estimates toward phylogenetic expectation, extreme fast-lived profiles may be dampened where trait data are sparse^49^, and host probability in these regions may consequently be underestimated. This gap removes observations as well as adding uncertainty: 7 species with confirmed positive detections were excluded from the models for lack of trait or phylogenetic data, including *Oligoryzomys utiaritensis* (nine positive individuals), *Microtus middendorffi* (seven) and *Micaelamys namaquensis* (four). All are Neotropical, Afrotropical or Siberian, and their exclusion is concentrated in the regions where trait coverage is weakest.

Our models project a static contemporary landscape, but as climate change redistributes mammalian biodiversity and drives agricultural expansion, the footprint of these interfaces will shift. Integrating this baseline biological trait framework with dynamic climate and land-use projections will be vital for tracking moving fronts of zoonotic hazard^50,51^. Moving from reactive outbreak response toward proactive, trait-guided monitoring is essential for anticipating emergence^52,53^.

## Methods

### Systematic Review & Data Harmonisation

We conducted a systematic review following PRISMA guidelines^54^ (Supplementary Figure S1). We searched PubMed and Web of Science for primary research published between January 1960, and August 2023, using search strings targeting *Arenaviridae* or *Hantaviridae* infection in small mammals (orders Rodentia, Eulipotyphla). Studies were included if they reported primary surveillance data (molecular or serological), identifiable host taxonomy and a geographic location for host detection. Data were extracted into a relational database capturing sampling location, sampling date, sampling effort, assay methodology, and number of individuals tested/positive^55^.

Host taxonomic harmonisation was performed using the taxize package (0.10) in R (4.2.3)^56,57^. All host names were resolved against the GBIF Backbone Taxonomy to handle synonyms and aligned with the Mammal Diversity Database (MDD v1.11) and IUCN Red List (v2024-2)^58–60^. Viral taxonomy was standardised to the International Committee on Taxonomy of Viruses (ICTV) 2024 release^61^. Viral taxonomic harmonisation was performed in Python using a family-aware reconciliation workflow with curated NCBI taxonomy lookup tables for *Arenaviridae* and *Hantaviridae*. Virus names were first normalised and matched to lookup records, prioritising species-level assignments. When exact species matches were unavailable, assignments were conservatively rolled up to higher lineage levels. Generic, placeholder, and ambiguous labels were handled with rule-based logic using genus context and metadata fields, and family-specific overrides were applied for known synonym or alias edge cases.

### Macroecological Trait Integration

Host life-history and ecological traits were aggregated from the COMBINE database^62^. We prioritised traits hypothesised to influence viral competence including adult mass (g), litter size (n), litters per year (n), gestation length (days), weaning age (days), maximum longevity (days) and age at first reproduction (days). Trait data were restricted to species in the target orders (Rodentia and Eulipotyphla). To maximise sample size without introducing list-wise deletion bias, we employed phylogenetic imputation using the Rphylopars (0.3.10) package^63^. This approach estimates missing trait values under a Brownian motion model of evolution, leveraging the phylogenetic covariance between species (using the consensus mammal tree) to predict missing values^64^.

Phylogenetic imputation produced one implausible value: maximum longevity for *Talpa altaica* was estimated at approximately 496 years. This species contributed no dyads to any model, having no recorded surveillance effort, and recomputing the principal component analysis with it excluded left PC1 loadings unchanged and scores correlated at *r* > 0.999. The value was retained without modification.

To resolve multicollinearity among life-history traits we performed a Principal Component Analysis (PCA) on log-transformed reproductive and morphological variables. The first principal component (PC1) explained 48.2% of the variance and was extracted as a continuous metric of the Pace of Life, where low values represent “fast” strategies (early maturity, high reproductive output) and high values represent “slow” strategies (high body mass, prolonged gestation and longevity).

### Quantification of Surveillance Bias

We assessed geographic bias using two complementary approaches. First, to quantify taxonomic coverage, we intersected species’ IUCN range maps with GADM (Level 2) administrative boundaries^65^. We aggregated these overlaps at the host genus level to calculate the proportion of each taxon’s geographic range that had been sampled, comparing the total area of surveyed districts to the full geographic range size of each genus. Second, to identify the sociodemographic drivers of surveillance intensity, we fitted a Zero-Inflated Negative Binomial (ZINB) model using the brms package (2.22)^66^. This model predicted the total count of tested individuals per district (aggregating all target species) as a function of night-time light intensity (VIIRS 2024)^67^, accessibility (travel time to major cities)^68^, and human population density^69^. The zero-inflation component modelled the probability of a district remaining entirely unsampled.

Local host species richness was derived by rasterising the filtered IUCN range polygons for the target mammalian orders onto the global predictor grid and summing the overlapping distributions to generate a continuous diversity surface.

Temporal trends were analysed using Generalised Additive Models (GAMs) with the mgcv package (1.9-3)^70^. We modelled the annual sampling effort (1960–2025) for each continent using a negative binomial error structure with thin plate regression splines to characterise non-linear trajectories and detect reactionary surveillance pulses following major outbreaks of zoonotic viruses.

To quantify the disconnect between field surveillance and genomic data availability, we compared the number of molecular detections (PCR or isolation) reported in the literature against the availability of sequence data in GenBank. Because primary literature frequently does not specify the exact subset of positive samples that successfully yielded viable sequences, we adopted a conservative overcounting approach: if a study reported molecular detections and linked corresponding sequences, we assumed all positive individuals from that specific record were sequenced. Using this conservative baseline, we calculated sequencing completeness ratios for both hosts and pathogens to identify regions where molecular detection is frequent but genomic data remains scarce.

### Phylogenetic Dyadic Modelling

To identify the intrinsic drivers of host status, we fitted a Bayesian Phylogenetic Dyadic Generalised Linear Mixed Model (GLMM). The unit of analysis was the host-virus dyad, defined as a potential association between a screened host species (*i*) and a viral species (*j*). To ensure that absence of a host-pathogen association reflected genuine negative data rather than a lack of surveillance, the analytic dataset was restricted to host species with at least one recorded testing event (Effort>0). Species present in the phylogeny but never sampled for *Arenaviridae* or *Hantaviridae* were excluded from the training set to avoid conflating absence of evidence with evidence of absence.

A binary indicator (*Y*_*ij*_) was coded as 1 if the dyad showed any evidence of natural viral interaction: active infection (PCR, sequencing, or viral isolation) or seroconversion (detectable antibodies reactive to the target virus). We acknowledge that seroconversion may reflect transient or dead-end infection rather than permissive or competent host status^71^; this primary model therefore maps the broad macroecology of detected host associations across the full breadth of historical surveillance. A secondary sensitivity analysis restricts this definition to active-infection evidence only. All other dyads within the sampled subset were set to 0 (pseudo-absences). The probability of host status (*p*_*ij*_ =*P*(*Y*_*ij*_ =1)) was thus modelled as:

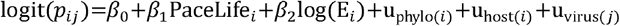

Where *log*(E_*i*_) is the log-transformed total count of individuals of host *i* tested for pathogens. While trap-nights provide a standardised measure of field effort, trap success and total trapping effort are inconsistently reported in historical literature, precluding their use without extensive and potentially circular imputation. By instead using the total number of individuals tested as a covariate, we capture the realised sampling volume. This controls for downstream biases: species that are rare, difficult to trap, or largely ignored yield fewer sampled individuals, resulting in lower statistical power, which the model accounts for by widening the respective uncertainty intervals. We acknowledge this effort measure may not reliably reflect family-specific testing intensity: surveillance is often geographically and epidemiologically targeted. Of individuals tested for these two families in the United States, 87.2% were tested for *Hantaviridae* versus 12.8% for *Arenaviridae*; in Guinea this was reversed 88.5% were tested for *Arenaviridae*, while testing in Nigeria and Sierra Leone was exclusively directed at *Arenaviridae* (100%). Therefore, a species with high total testing volume in one region may still be comparatively under-tested for the locally non-prioritised viral family.

The model was applied to three analytic datasets:

1. The Global Model: Comprising 43,677 dyadic pairs from 69 pathogens (including unclassified *Arenaviridae* and *Hantaviridae*) and 633 rodent and shrew species. In this model, we tested the life-history hypothesis using pace of life (PC1) as the primary predictor. This model was subsequently used for global spatial projection to maximise taxonomic coverage.
2. The Synanthropy Subset: Comprising 18,906 dyadic pairs containing the same 69 pathogens but a reduced set of 274 host species for which detailed synanthropy data were available. In this subset, the life-history and ecological-opportunity hypotheses were tested in tandem through the addition of a categorical synanthropy term (Not Synanthropic, Occasionally Synanthropic, Totally Synanthropic)^22^. This model was used to test whether synanthropy predicts host status independently of pace of life.
3. The Complete-Case Subset: Comprising 14,697 dyadic pairs from the same 69 pathogens but a restricted dataset of 213 host species possessing entirely empirical, non-imputed life-history trait values. This subset was used strictly to verify that macroecological signals were not artefacts of phylogenetic imputation.

Of the 637 species assayed for these two viral families, 542 entered the global model. Of those excluded, 65 fell outside the target orders and 30 lacked the trait or phylogenetic data required for model fitting. The model additionally includes 91 species that were sampled and tested for other pathogens but not for these two families; because sampling effort is summed across all pathogens, these species contribute dyads with recorded effort and no positive detections.

We incorporated three random effects to account for non-independence. u_phylo(*i*)_ is a structured random phylogenetic effect covarying according to the phylogenetic distance matrix (*A*) derived from the consensus tree, controlling for shared evolutionary history^64^. u_host(*i*)_ is an unstructured species-specific random intercept included to capture species-level variation (e.g., overdispersion) not explained by phylogeny. Finally, viral identity u_virus(*j*)_ is included as a random intercept controlling for uneven study effort across viral species (e.g., *Orthohantavirus sinnombreense* is studied more intensely than South American hantaviruses).

To ensure our macroecological inferences were not confounded by historical serological cross-reactivity, we conducted sensitivity analyses. First, we subset the global dyadic data to include only high-confidence host assignments, redefining *Y*_*ij*_ =1 strictly as dyads with molecular evidence (PCR or sequencing) or viral isolation. The Bayesian GLMM was re-fitted to this restricted dataset using identical priors and chain parameters to assess the stability of the pace of life (PC1) coefficient. Second, to verify that the observed trait associations were not artefacts of phylogenetic imputation, we conducted a complete-case sensitivity analysis. The model was refitted to the Complete-Case Subset (*n* = 213), with the PCA and phylogenetic covariance matrices recomputed strictly for these empirical values. We adopted a Bayesian framework to ensure our estimates reflect ecologically meaningful effect sizes with propagated uncertainty, rather than relying on null-hypothesis significance testing, which is prone to *p*-value inflation given our substantial sample size.

Models were fitted in Stan via the brms interface using 8 chains of 2,500 iterations, including 1,500 warmup iterations. We specified weakly informative priors to regularise parameter estimates (Normal(0,1) for fixed effects; Exponential(1) for variance components). Convergence was assessed via 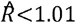 and visual inspection of trace plots.

### Global Host Probability Mapping

To validate the out-of-sample predictive power of our spatial projections, we first executed a Leave-One-Region-Out (LORO) spatial cross-validation. The dataset was partitioned by continent, and the Bayesian GLMM was iteratively refitted, withholding one continent’s data per iteration. Out-of-sample host probabilities were predicted for the withheld regions based exclusively on the biological relationships learned from the remaining global data. Species were assigned to regions and to cross-validation folds by majority occurrence. Each species was allocated to the region, or continent, containing the majority of its recorded country-level occurrences. Species whose occurrences span multiple regions are thus assigned to a single region, preserving independence between the groups compared. Western Europe was excluded from the regional comparison, as only three species met the majority-occurrence criterion.

We assessed predictive performance in two ways. First, we scored predictions against observed host status (*Y*_*ij*_) for all withheld dyads using the area under the receiver operating characteristic curve (AUC), evaluating predictions at each dyad’s actual recorded sampling effort rather than a fixed reference value, since observed host status is itself conditional on realised effort. Second, we compared out-of-sample predictions against full in-sample model estimates using Spearman’s rank correlation, assessing coefficient stability across folds. Because species are not randomly distributed across continents, we additionally quantified genus-level overlap between each withheld fold and its training set, as shared genera reduce the extent to which withheld regions represent genuinely novel biology.

Following validation, we mapped the final global distribution of community host probability by projecting the parameters of the Global Model (trained on Life History and Sampling Effort) onto the full set of 2,766 mammal species for which life history data were available. We generated posterior predictions for host probability (*p*_*i*_) for all species, setting sampling effort to the 95th percentile of observed values to estimate biological potential independent of surveillance history. We used the 95th percentile rather than the observed maximum because the maximum corresponds to a single species (*Peromyscus maniculatus*), itself a product of the surveillance patterns described above; predicted probabilities differ by approximately eight-fold between the two reference points.

Spatial projection was performed using the terra package (1.8-60)^72^. We rasterised IUCN Red List range maps for all 2,766 species onto a Mollweide equal-area projection at 20km resolution. A 20km resolution was selected to balance global computational tractability against the spatial precision of IUCN range polygons. For each grid cell (*c*), we calculated Community Host Probability (*H*_*c*_) as the arithmetic mean of the predicted host probabilities for all species present in that cell:

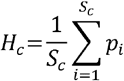

Where *S*_*c*_ is the species richness in cell *c*. To avoid artefacts from species-poor pixels, cells with *S*_*c*_ <5 were masked. We note that *H*_*c*_ represents the host-side component of intrinsic zoonotic hazard only: viral random effects were marginalised in generating *p*_*i*_, so *H*_*c*_ does notincorporate pathogen-specific factors (virulence, transmissibility, or human severity) that jointly determine realised hazard alongside host community composition.

To explicitly interrogate the relationship between biodiversity and community host probability, we performed a bivariate classification using the biscale package (1.1)^73^. Grid cells were categorised into a 3×3 matrix based on tertiles (33rd and 66th percentiles) of species richness and community host probability. We extracted mean values for all administrative districts (GADM Level 2) using the exactextractr package (0.10)^74^. Because area-weighted extraction ignores masked cells and renormalises over the remainder, districts with limited coverage receive values derived from a fraction of their area. We therefore retained only districts for which valid cells covered at least half the district (37,676 of 38,367; 1.8% excluded). Excluded districts are predominantly small island and coastal regions, where a 20km cell frequently spans both land and sea.

### Analysis of Evolutionary Congruence

To test evolutionary congruence between viruses and their mammal hosts, we analysed *Arenaviridae* and *Hantaviridae* separately using binary host-virus association matrices derived from harmonised surveillance records. Associations were restricted to PCR-positive detections, meaning assays reporting at least one positive individual by PCR, excluding serological assays, so that each link represents evidence of active infection. Broad and unresolved taxa (e.g., family-level and unclassified labels) were removed prior to matrix construction. For each family, we retained only observed links by transforming interaction counts into binary incidence, then pruned matrices to taxa shared between association data and phylogenies (dropping empty rows and columns after pruning). Host phylogenies were taken from the pruned VertLife mammal tree^64^; viral phylogenies were built from family-specific nucleoprotein HIPSTR trees^75^ inferred using BEAST X^76^ under a coalescent with constant population model and a strict clock.

Cophylogenetic signal was quantified with the Procrustean Approach to Cophylogeny (PACo)^77,78^. Briefly, cophenetic distance matrices were computed for host and viral trees, assembled with the binary association matrix, and projected into a principal-coordinate space with Cailliez correction to handle potential non-Euclidean structure. PACo superimposition was run with 999 permutations, treating host phylogeny as the reference configuration and fitting viral phylogeny onto it. Global congruence was evaluated with the PACo goodness-of-fit statistic 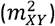; in this framework, a lower residual contribution indicates stronger host-virus phylogenetic congruence for a given link.

As a complementary global-fit test, we also ran ParaFit on the same host and viral distance matrices and association matrix. PACo link-level residuals were extracted and exported per host-virus pair for downstream analyses of residual distributions and visualisation.

### Host Phylogenetic Diversity Accumulation and Saturation Analysis

We quantified the realised host breadth for each viral species using a branch-length-based host phylogenetic diversity (PD) framework on the pruned mammalian host phylogeny. Viral identity was defined as previously described, and broad or unresolved labels were excluded before analysis. Host-virus records were standardised to phylogeny-compatible host tip labels, and only associations with resolvable hosts and valid year metadata were retained. Associations were restricted to infected hosts, defined by a positive PCR, sequencing or virus culture result. For each host-virus pair, the first detection year was defined as the earliest year observed, producing a virus-specific temporal sequence of unique host discoveries.

Faith’s PD was computed as the sum of branch lengths in the minimal subtree spanning the accumulated host set^79^. For each virus, cumulative PD (PD _*c*_) was calculated in two ways: (i) year-batched accumulation, in which all hosts first detected in the same year were added together, and (ii) host-ordered accumulation, in which hosts were added sequentially by first detection year (with deterministic tie handling). At each step (*t*), we calculated cumulative PD and incremental gain (*ΔPD*_*t*_). We then summarised per-virus host breadth using final host richness, final PD, normalised PD, PD per host, maximum increment, and relative burstiness. To characterise host phylogenetic dispersion independently of PD, we additionally estimated mean pairwise patristic distance (MPD) and mean nearest taxon distance (MNTD) among occupied hosts.

To test whether observed host-order accumulation deviated from random expectation, we generated permutation null envelopes by randomising host discovery order without replacement and recomputing cumulative PD. In per-virus analyses, permutations were performed on each virus’s observed host set. At each host index, we summarised the null as the permutation mean and the 2.5–97.5% interval. For family-level comparisons (*Arenaviridae* vs *Hantaviridae*), hosts were pooled within each family and ordered by first detection year. Cumulative PD was computed on the raw host-index scale (cumulative discovered hosts), without axis normalisation. Family-level null expectations were then generated by randomising discovery order within each pooled family host set (10,000 permutations) and extracting the 2.5–97.5% envelope at each host index. We tested whether discovery order was phylogenetically clustered by comparing the area under the observed family-level cumulative PD curve with the same statistic across the 10,000 permutations (one-tailed test). Ties among hosts first detected in the same year were broken at random, and the statistic was averaged over 1,000 tie-breaks.

## Acknowledgements

This research was supported by funding to Verena (viralemergence.org) from the U.S. National Science Foundation (NSF DBI 2515340 to S.N.S, supporting R.R.). DS is supported by a Fellowship-in-Residence from Verena and receives salary support from the NSF-NIH-NIFA Ecology and Evolution of Infectious Disease Award #2208034 in partnership with Research and Innovation (UKRI) Biotechnology and Biological Sciences Research Council (BBSRC) Award BB/X005364/1 (awarded to DWR). S.N.S. is partly supported by the Centers for Disease Control and Prevention Center for Forecasting and Outbreak Analytics (cooperative agreement CDC-RFA-FT-23-0069). R.R. was partly supported by the Poncin Scholarship Fund from the Poncin Trust under award number GF007810.

## Author Contributions

**Conceptualisation:** DS, RR, SNS; **Methodology:** DS, RR, SNS, DWR; **Software:** DS, RR; **Formal Analysis:** DS, RR, SNS, DWR; **Resources:** DS; **Data Curation:** DS, RR, GR, AM-C, HG; **Writing – Original Draft:** DS, RR; **Writing – Review & Editing:** DS, RR, GR, AM-C, HG, DWR, SNS; **Visualisation:** DS, RR; **Supervision:** DS, SNS, DWR; **Project Administration:** DS, SNS, DWR; **Funding Acquisition:** DS, SNS, DWR.

## Data Availability

All analytical code, computational pipelines, and the primary ArHa database required to reproduce these analyses are accessible at https://github.com/DidDrog11/arha-macroecology and archived on Zenodo https://doi.org/10.5281/zenodo.20523966, in accordance with FAIR data principles. Due to file size constraints and guidance from data providers, the repository does not host the environmental spatial layers or IUCN range polygons. Instead, the reproducible R scripts required to fetch, filter, and harmonise the remaining spatial and ecological data directly from their original public sources are documented.

## Statement on AI use

Generative AI tools were used in the preparation of this manuscript: Google Gemini (3.1 Pro) for language and structure of the main text, and Anthropic Claude (Opus 5.5) for code review and documentation, consistency checking between outputs and reported values. No AI tool generated data, conducted analyses, or made interpretive decisions. DS reviewed and verified all AI-assisted text and code, and all authors take full responsibility for the content.

