## Supplemental File for "Fast pace of life is associated with host status in small mammals, despite pervasive surveillance bias"

### Supplementary Material

#### The Genetic Data Gap

Beyond the biases in host sampling, we identified a substantial disconnect between viral detection and genomic characterisation. Of the 8,063 individual hosts that tested positive via molecular assays or isolation, valid sequence data was available for only 41.4% (3,340 individuals). Sequencing completeness varied by viral family: *Hantaviridae* detections had lower genomic coverage (38.1%) compared to *Arenaviridae* (55.5%). Spatially, the proportion of unsequenced positive results was highest in Northern America (84.1%) (Supplementary Figure S9). As a result, 67.2% of the unique host-virus associations in the global dataset are currently supported only by serological or fragment-based evidence without lineage-specific sequence confirmation (Supplementary Figure S10).

#### Host Discovery Trajectories for Exemplar Viruses

To provide mechanistic context for the accumulation of host phylogenetic diversity ($PD_{c}$), we tracked the discovery trajectories of six well-sampled viral exemplars: *Orthohantavirus andesense*, *O. hantanense*, *O. seoulense*, *O. sinnombreense*, *O. tulaense*, and *Mammarenavirus choriomeningitidis* (Supplementary Figure S8). All accumulated host phylogenetic diversity in discrete steps rather than smoothly, but the events behind those steps differ between viruses, and only some align with recognised human outbreaks.

The clearest outbreak-driven trajectory is *O. sinnombreense*. A single host was recorded in 1993, followed by six further hosts in 1994 contributing the largest single-year increment in the dataset ($\Delta PD_{t}$ = 241.3). This aligns with the 1993 Four Corners hantavirus pulmonary syndrome emergence and the intensification of follow-up surveillance that accompanied it^41^. For *M. choriomeningitidis* (LCMV), the earliest recorded increment follows the 2005 United States transplant-associated cluster^80^, although the largest increments in its trajectory are recent (2008, 2021 and 2022), reflecting renewed attention to this virus rather than a single spillover event.

For the remaining exemplars we did not identify clear links to single, large, recognised human outbreaks. *O. tulaense* accumulated most of its host phylogenetic diversity in one step in 2014, with smaller later increments, a pattern more consistent with rodent-focused field campaigns and the regional expansion of molecular surveillance than with reactive case investigation^81^. *O. hantanense* and *O. seoulense* likewise accumulated hosts in a small number of widely separated steps, with the largest increments in 1995 and 1999 for *O. hantanense* and in 2004 and 2012 for *O. seoulense*. For both viruses the largest increments were produced by detections outside the rodent hosts with which they are usually associated, principally in shrews (*Suncus murinus*, *Crocidura suaveolens*, *Crocidura lasiura* and *Sorex araneus*), which lie in a different mammalian order and therefore contribute disproportionate branch length. The studies reporting these detections were surveys of wider host communities rather than investigations of human cases, including an explicit search for reservoir host expansion^82^ and a molecular survey of hantaviruses across co-occurring rodent species^83^. *O. andesense* accumulated hosts steadily between 2000 and 2009, with one substantial increment in 2009 and no further additions after 2014.

Taken together these trajectories support the family-level result rather than contradicting it. Host discovery proceeds in steps for every exemplar, but those steps are driven by reactive case investigation for only a subset of viruses, and the phylogenetic clustering of discovery order reported in the main text is the more general pattern.

#### Supplementary Model Summaries

Table S1 details the posterior summaries and MCMC diagnostics for the Zero-Inflated Negative Binomial (ZINB) spatial surveillance model. This model was constructed to disentangle the socioeconomic drivers of global sampling intensity from underlying host biodiversity at the administrative district level. The parameter estimates are partitioned into two distinct processes: the count model, which estimates the magnitude of sampling effort in surveyed areas, and the zero-inflation model, which predicts the probability of a district remaining entirely unsampled.

Consistent with a pervasive “streetlight effect,” proxies for anthropogenic development and logistical convenience dominate. Night-time light intensity predicts both a higher volume of testing (count) and a reduced probability of remaining unsampled (zero-inflation). Conversely, local host species richness exhibits no credible effect on the number of individuals tested (95% CrI [−0.271, 0.110]), consistent with global surveillance being associated with human infrastructure rather than ecological or biological relevance. MCMC diagnostics ($\hat{R}\approx1.00$ and high Effective Sample Sizes) indicate robust convergence across all parameters.

| Table S1 \| Posterior summary and MCMC diagnostics for the Zero-Inflated Negative Binomial (ZINB) spatial surveillance model. | | | | | | | |
| --- | --- | --- | --- | --- | --- | --- | --- |
| Parameter | Posterior Mean | SE | 2.5% CrI | 97.5% CrI | R̂ | Bulk ESS | Tail ESS |
| **Fixed Effects** | | | | | | | |
| Intercept (Count) | -5.029 | 0.245 | -5.504 | -4.540 | 1.005 | 675.5957 | 1149.5120 |
| Intercept (Zero-Inflation) | -5.041 | 1.785 | -8.807 | -1.531 | 1.013 | 581.8182 | 434.7106 |
| Population Density (Count) | 0.215 | 0.165 | -0.111 | 0.529 | 1.000 | 1834.6812 | 2512.6319 |
| Night-light Intensity (Count) | 0.501 | 0.174 | 0.171 | 0.853 | 1.000 | 1620.7501 | 1933.0036 |
| Accessibility (Count) | -0.485 | 0.150 | -0.767 | -0.184 | 1.000 | 1814.7728 | 2252.3709 |
| Host Richness (Count) | -0.088 | 0.097 | -0.271 | 0.110 | 1.000 | 2339.3998 | 2875.6937 |
| Population Density (Zero-Inflation) | 1.040 | 0.532 | 0.116 | 2.152 | 1.005 | 1334.0278 | 1598.3460 |
| Night-light Intensity (Zero-Inflation) | -5.076 | 2.108 | -9.522 | -1.272 | 1.011 | 659.0640 | 515.9400 |
| Accessibility (Zero-Inflation) | -1.991 | 0.698 | -3.541 | -0.837 | 1.008 | 776.1669 | 1103.5836 |
| Host Richness (Zero-Inflation) | -0.684 | 0.317 | -1.361 | -0.103 | 1.008 | 851.5409 | 824.6626 |
| **Variance Components** | | | | | | | |
| σ (Country / GID_0) | 1.899 | 0.195 | 1.542 | 2.306 | 1.002 | 942.3438 | 1601.8107 |
| **Distributional Parameters** | | | | | | | |
| Shape (Negative Binomial) | 0.007 | 0.000 | 0.006 | 0.008 | 1.008 | 793.1948 | 446.0273 |

Table S2 details the posterior summaries and MCMC diagnostics for the Global Phylogenetic Dyadic GLMM ($N$ = 43,677). This model tests the macroecological association between a host’s pace of life (PC1) and its host status, whilst controlling for sampling volume (Log(Sampling Effort)) and non-independence via random effects. The model demonstrates that sampling effort is the dominant predictor of observed host status. After adjusting for this effort, a fast pace of life (lower PC1 scores) exhibits a negative posterior mean, indicating higher host probability, though the 95% credible interval crosses zero. Variance partitioning reveals that host status is a highly labile, species-specific trait ($\sigma_{Host}\text{=}0.418$), outweighing deep evolutionary conservation ($\sigma_{Phylogeny}\text{=}0.098$).

| Table S2 \| Posterior summary and MCMC diagnostics for the Full Dyadic Model (N = 43,677). Variance components are reported as standard deviations (σ). | | | | | | | |
| --- | --- | --- | --- | --- | --- | --- | --- |
| Parameter | Posterior Mean | SE | 2.5% CrI | 97.5% CrI | R̂ | Bulk ESS | Tail ESS |
| **Fixed Effects** | | | | | | | |
| Intercept | -8.194 | 0.890 | -9.904 | -6.365 | 1.000 | 5916.742 | 5247.209 |
| Log(Sampling Effort) | 0.585 | 0.029 | 0.530 | 0.644 | 1.000 | 6955.780 | 6323.804 |
| Pace of Life (PC1) | -0.125 | 0.084 | -0.289 | 0.037 | 1.000 | 9372.240 | 6051.342 |
| **Variance Components** | | | | | | | |
| σ (Host Species) | 0.418 | 0.106 | 0.183 | 0.607 | 1.004 | 1671.158 | 1934.578 |
| σ (Viral Species) | 1.488 | 0.150 | 1.220 | 1.809 | 1.003 | 2074.891 | 4388.244 |
| σ (Phylogeny) | 0.098 | 0.019 | 0.063 | 0.136 | 1.002 | 2307.763 | 4340.714 |

Table S3 presents the subset analysis ($N$ = 18,906) evaluating the ecological opportunity hypothesis by incorporating a categorical synanthropy term alongside life-history traits. The model provides some support that synanthropy acts as a distinct predictor of host probability, independent of life history. Totally synanthropic (obligate commensal) species exhibit a moderate positive association with host status (Posterior Mean = 0.762, CrI = 0.045-1.46) relative to non-synanthropic species. The inclusion of human affinity does not diminish the biological signal of life history; the PC1 coefficient remains stable (Posterior Mean = -0.147).

| Table S3 \| Posterior summary and MCMC diagnostics for the Synanthropy Subset Model (N = 18,906). | | | | | | | |
| --- | --- | --- | --- | --- | --- | --- | --- |
| Parameter | Posterior Mean | SE | 2.5% CrI | 97.5% CrI | R̂ | Bulk ESS | Tail ESS |
| **Fixed Effects** | | | | | | | |
| Intercept | -7.969 | 0.952 | -9.827 | -6.082 | 1.001 | 6435.366 | 5246.074 |
| Log(Sampling Effort) | 0.450 | 0.039 | 0.376 | 0.530 | 1.001 | 6527.350 | 5277.290 |
| synanthropy_statusOccasionallySynanthropic | 0.232 | 0.157 | -0.075 | 0.540 | 1.001 | 8037.261 | 6470.341 |
| synanthropy_statusTotallySynanthropic | 0.762 | 0.359 | 0.045 | 1.456 | 1.000 | 7227.072 | 5883.264 |
| Pace of Life (PC1) | -0.147 | 0.091 | -0.332 | 0.025 | 1.000 | 8199.437 | 6413.444 |
| **Variance Components** | | | | | | | |
| σ (Host Species) | 0.293 | 0.138 | 0.029 | 0.546 | 1.004 | 1258.398 | 2365.334 |
| σ (Viral Species) | 1.854 | 0.211 | 1.483 | 2.304 | 1.003 | 2266.808 | 3805.997 |
| σ (Phylogeny) | 0.097 | 0.021 | 0.059 | 0.140 | 1.004 | 2164.322 | 4084.023 |

Table S4 details the strict sensitivity analysis, which restricts positive host status exclusively to molecular detection and viral isolation to remove potential historical serological cross-reactivity and dead-end spillover noise. By isolating active viral infection, the biological signal linking a fast pace of life to host probability remains consistent (Posterior Mean = -0.167, compared to -0.125 in the global model). Furthermore, restricting the definition to molecular evidence appropriately reduces the variance attributed to specific viral species ($\sigma_{Viral}$ drops to 1.251 from 1.488) whilst increasing idiosyncratic host-level variance ($\sigma_{Host}$ rises to 0.610), suggesting that historical serological datasets introduce homogenising noise across diverse host species.

| Table S4 \| Posterior summary and MCMC diagnostics for the Strict Molecular Sensitivity Model. | | | | | | | |
| --- | --- | --- | --- | --- | --- | --- | --- |
| Parameter | Posterior Mean | SE | 2.5% CrI | 97.5% CrI | R̂ | Bulk ESS | Tail ESS |
| **Fixed Effects** | | | | | | | |
| Intercept | -8.356 | 1.107 | -10.461 | -6.089 | 1.000 | 6559.728 | 6157.653 |
| Log(Sampling Effort) | 0.600 | 0.043 | 0.520 | 0.689 | 1.001 | 7733.218 | 6194.514 |
| Pace of Life (PC1) | -0.167 | 0.120 | -0.400 | 0.066 | 1.001 | 8898.171 | 5773.085 |
| **Variance Components** | | | | | | | |
| σ (Host Species) | 0.610 | 0.158 | 0.251 | 0.896 | 1.004 | 1646.994 | 1694.400 |
| σ (Viral Species) | 1.251 | 0.148 | 0.990 | 1.573 | 1.001 | 3170.119 | 4550.569 |
| σ (Phylogeny) | 0.132 | 0.032 | 0.075 | 0.200 | 1.002 | 1526.469 | 2760.181 |

Table S5 details the complete-case empirical sensitivity analysis ($N$ = 14,697). To ensure the observed macroecological signals are not mathematical artefacts of phylogenetic trait imputation, this model restricts the dataset strictly to host species possessing entirely empirical, non-imputed values for all life-history variables. The posterior mean for the empirical Pace of Life (PC1) remains stable (-0.148), aligning closely with the full imputed model. Variance partitioning within this strict subset confirms that species-specific variation ($\sigma_{Host}\text{=}0.358$) continues to outweigh phylogenetic covariance ($\sigma_{Phylogeny}\text{=}0.091$) in the absence of an imputed phylogenetic structure.

| Table S5 \| Posterior summary and MCMC diagnostics for the Complete-Case Empirical Model (N = 14,697). | | | | | | | |
| --- | --- | --- | --- | --- | --- | --- | --- |
| Parameter | Posterior Mean | SE | 2.5% CrI | 97.5% CrI | R̂ | Bulk ESS | Tail ESS |
| **Fixed Effects** | | | | | | | |
| Intercept | -8.733 | 0.908 | -10.485 | -6.890 | 1.002 | 4048.1305 | 4630.740 |
| Log(Sampling Effort) | 0.617 | 0.044 | 0.537 | 0.707 | 1.000 | 5280.2137 | 5623.306 |
| Pace of Life (PC1 - Empirical) | -0.148 | 0.111 | -0.374 | 0.064 | 1.000 | 5626.3393 | 5946.245 |
| **Variance Components** | | | | | | | |
| σ (Host Species) | 0.358 | 0.155 | 0.046 | 0.645 | 1.016 | 929.6325 | 1597.169 |
| σ (Viral Species) | 1.931 | 0.234 | 1.522 | 2.430 | 1.006 | 1513.9528 | 3555.603 |
| σ (Phylogeny) | 0.091 | 0.027 | 0.043 | 0.146 | 1.009 | 1354.0717 | 2480.014 |

Table S6 provides definitions, units, and empirical distributions for the seven life-history traits used to construct the Pace of Life axis (PC1). The proportion of values obtained by phylogenetic imputation rather than direct measurement is reported for each trait, as trait coverage varies substantially: adult body mass is recorded for the large majority of species, whereas reproductive timing traits are missing for the majority and are correspondingly reliant on imputation. Summary statistics are calculated from empirically recorded values only, prior to imputation.

Table S6 | Life-history covariate definitions and empirical distributions. Summary statistics reflect empirically recorded values only; % imputed indicates the proportion of species for which the value was estimated by phylogenetic imputation. Distributions are shown on a log₁₀ scale.

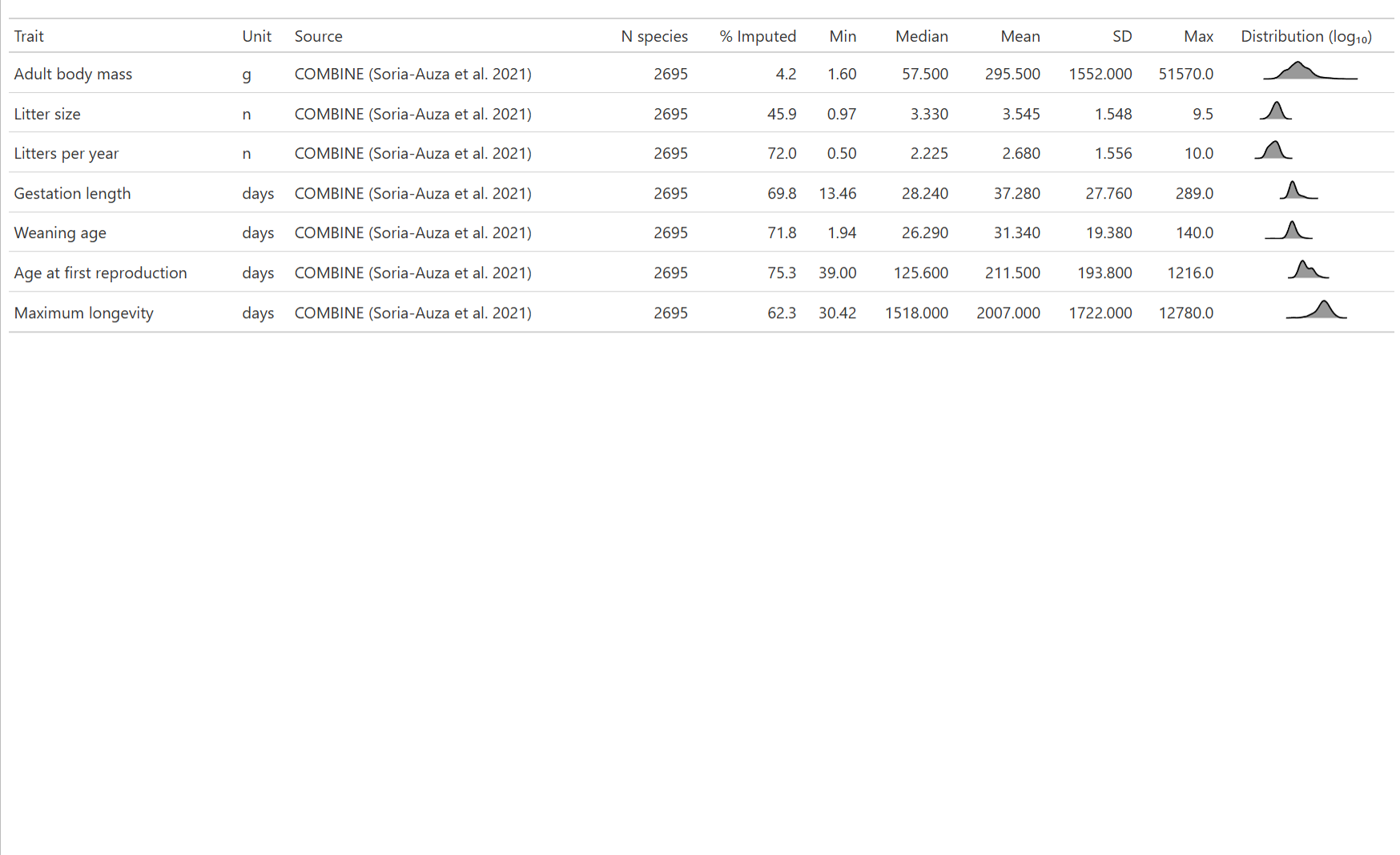

Table S7 provides equivalent definitions and distributions for the district-level covariates used in the Zero-Inflated Negative Binomial model of surveillance intensity. Values are reported in native units; all covariates were standardised prior to model fitting.

| Table S7 \| Spatial covariate definitions and empirical distributions for the ZINB surveillance model (N = 38,897 administrative districts). | | | | | | | | |
| --- | --- | --- | --- | --- | --- | --- | --- | --- |
| Covariate | Unit | Source | N districts | Min | Median | Mean | SD | Max |
| Host species richness | mean species count (raster mean) | IUCN range rasterisation | 38897 | 0 | 20.1700 | 19.790 | 7.378 | 55.82 |
| Night-time light intensity | VIIRS radiance (nW/cm²/sr) | VIIRS 2024 | 38897 | 0 | 0.4525 | 1.906 | 5.596 | 236.70 |
| Population density | people / km² | WorldPop 2025 | 38897 | 0 | 62.1800 | 307.100 | 1073.000 | 20760.00 |
| Accessibility (travel time to nearest city) | minutes | Weiss et al. 2018 | 38897 | 0 | 29.0500 | 69.230 | 195.100 | 11760.00 |

Table S8 disaggregates the genetic data gap by geographic region. Sequencing completeness varies more than six-fold across regions, with Northern America showing the largest disparity between molecular detection and available sequence data. Denominators are pathogen-positive individuals detected by PCR, sequencing, or isolation; regions with fewer than 50 positive individuals are excluded.

| Table S8 \| Sequencing completeness by geographic region. Denominator is individuals testing positive by molecular assay or isolation. Regions with fewer than 50 positive individuals are excluded. | | | | |
| --- | --- | --- | --- | --- |
| Region | Positive hosts | Sequenced | % Sequenced | % Unsequenced |
| Northern America | 1436 | 229 | 15.9 | 84.1 |
| Southern Europe | 262 | 83 | 31.7 | 68.3 |
| Western Europe | 1546 | 633 | 40.9 | 59.1 |
| Eastern Asia | 1974 | 895 | 45.3 | 54.7 |
| Eastern Europe | 349 | 171 | 49.0 | 51.0 |
| Sub-Saharan Africa | 1197 | 645 | 53.9 | 46.1 |
| Latin America and the Caribbean | 543 | 318 | 58.6 | 41.4 |
| Northern Europe | 293 | 257 | 87.7 | 12.3 |

Table S9 reports out-of-sample performance for the Leave-One-Region-Out cross-validation. Discrimination was assessed by AUC against observed host status at each dyad’s recorded sampling effort. Genus-level overlap between each withheld fold and its corresponding training set is reported alongside, as shared genera reduce the extent to which a withheld continent represents biologically novel taxa.

| Table S9 \| Leave-One-Region-Out cross-validation performance and phylogenetic overlap between withheld and training folds. | | | | | | |
| --- | --- | --- | --- | --- | --- | --- |
| Withheld continent | AUC | N dyads | N positive | Genera (withheld) | Shared with training | % Shared |
| Americas | 0.810 | 20010 | 324 | 91 | 8 | 8.8 |
| Asia | 0.832 | 9246 | 93 | 60 | 16 | 26.7 |
| Europe | 0.826 | 3519 | 95 | 24 | 9 | 37.5 |
| Africa | 0.876 | 9936 | 55 | 54 | 7 | 13.0 |

### Supplemental Figures

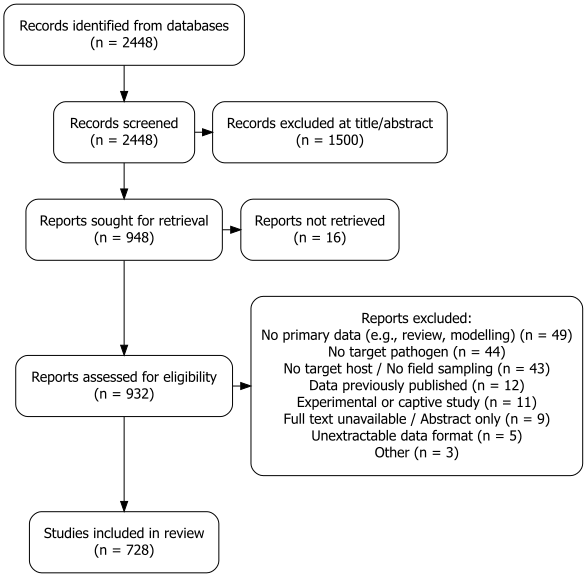
**Supplementary Figure 1 | Systematic review and data harmonisation workflow.** PRISMA flow diagram detailing the literature search, screening, and inclusion process for primary surveillance records of *Arenaviridae* and *Hantaviridae* in small mammals.

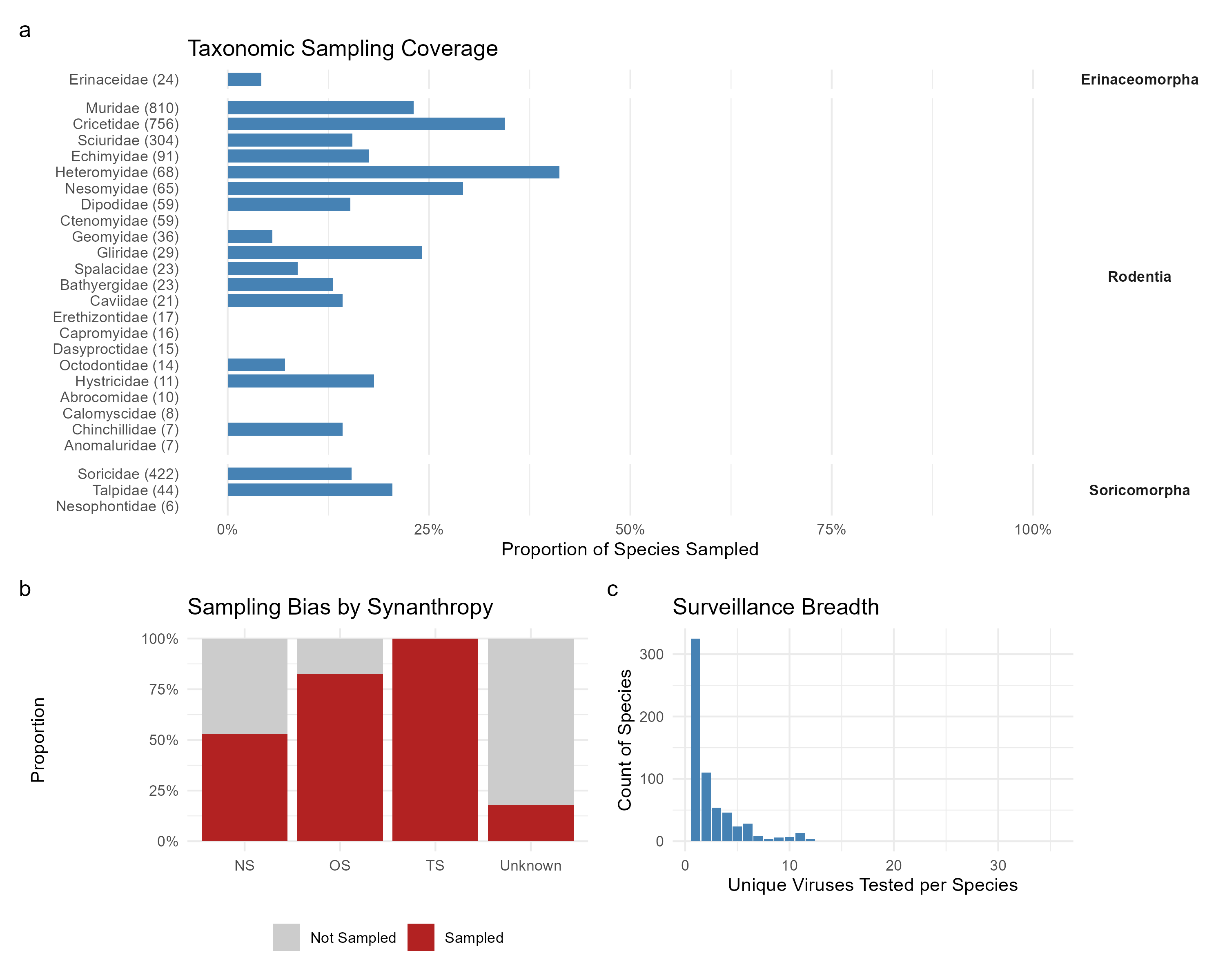

**Supplementary Figure 2 | Taxonomic sampling bias and surveillance breadth.** a) Proportion of sampled versus unsampled species across mammal families within the target orders, highlighting complete surveillance gaps in families such as *Gliridae*. b) Sampling bias by synanthropy status, showing the disproportionate sampling of synanthropic species relative to their abundance in the global database. c) Surveillance breadth, illustrating that the vast majority of sampled host species are tested for only a single virus.

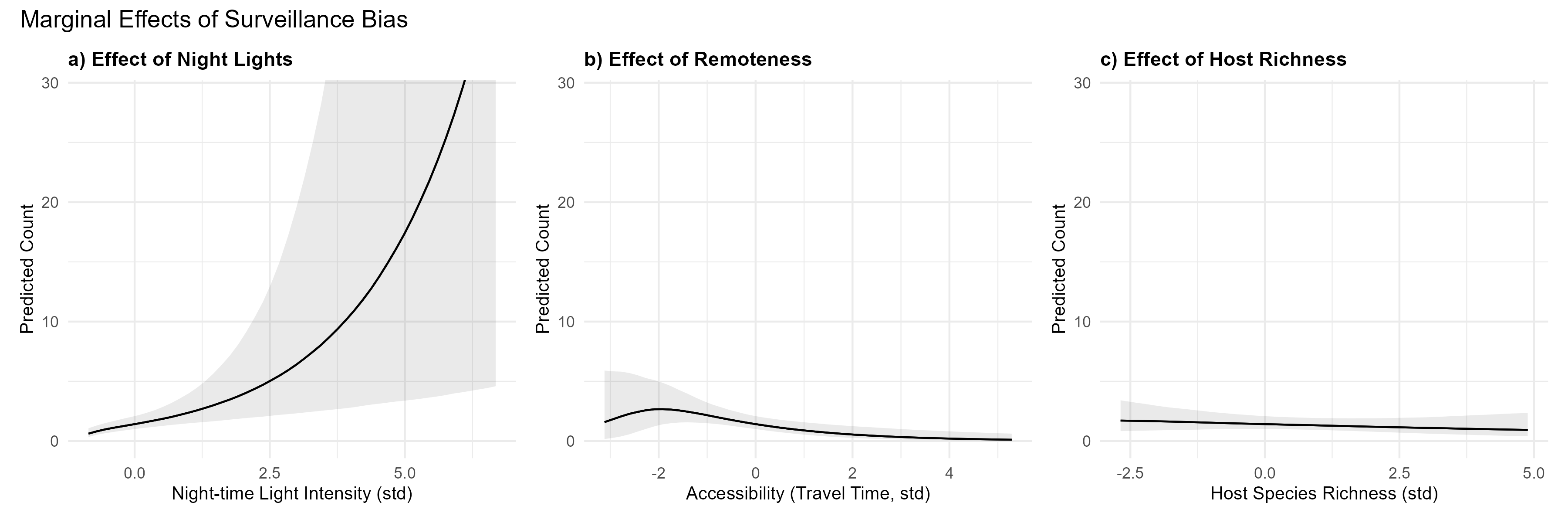

**Supplementary Figure 3 | Marginal effects of sociodemographic drivers on surveillance effort.** Predicted sampling intensity (number of individuals tested) as a function of a) night-time light intensity, b) accessibility (travel time to major cities), and c) local host species richness, derived from the Zero-Inflated Negative Binomial (ZINB) spatial model. Surveillance effort increases with infrastructure and accessibility but is decoupled from local biodiversity.

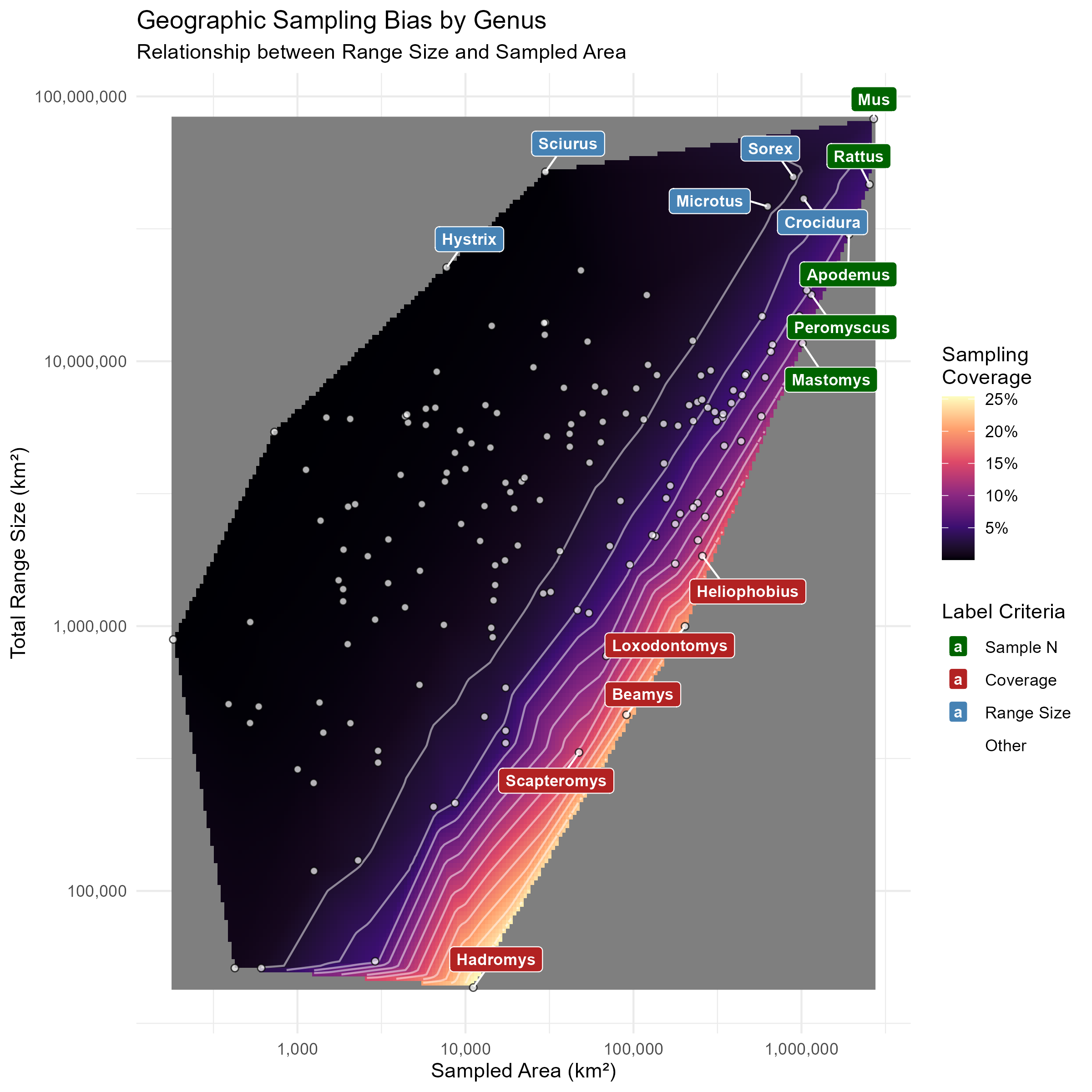

**Supplementary Figure 4 | The disconnect between geographic distribution and surveillance coverage.** The relationship between the total geographic range size of a small mammal genus (y-axis) and the total area of administrative districts in which it has been sampled for *Arenaviridae* or *Hantaviridae* (x-axis). Both axes are log-transformed. Points represent individual genera ($N\text{=}186$). The background heatmap and contours visualise the proportional coverage (Sampled Area / Total Range), interpolated from the observed data, with lighter colours indicating higher coverage. Labels highlight genera that represent extremes in total sample size (green), proportional coverage (red), or geographic range size (blue). Despite a positive correlation between range size and sampled area, the vast majority of genera occupy the dark purple region, indicating that even widespread taxa are sampled across only a small fraction of their distribution.

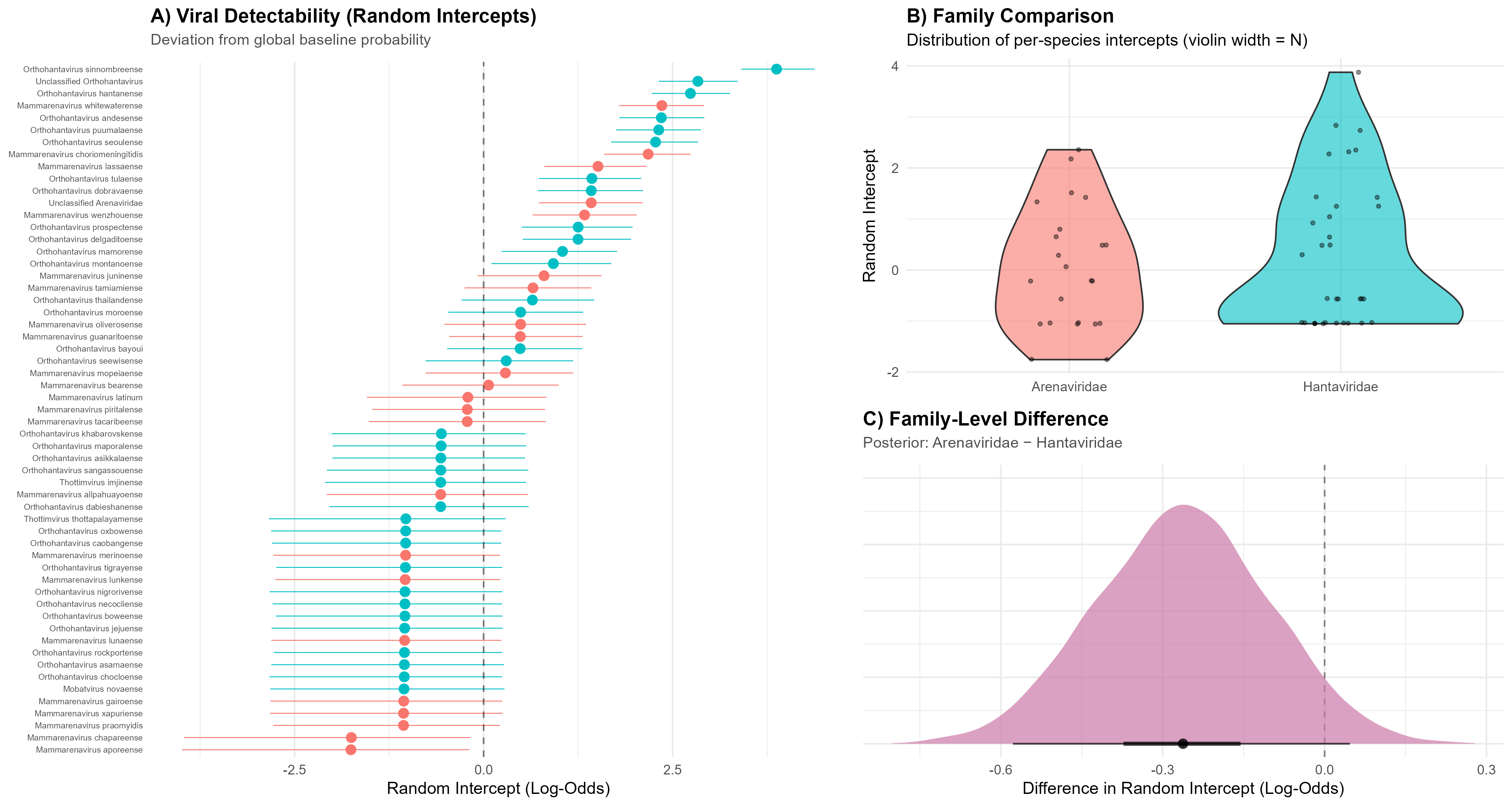

**Supplementary Figure 5 | Viral detectability by species and family.** a) Caterpillar plot of random intercepts (log-odds) for individual viral species, representing their deviation from the global baseline probability of detection in the Bayesian GLMM; points show posterior medians with 95% credible intervals. b) Violin plot comparing the distribution of per-species random intercepts between *Arenaviridae* and *Hantaviridae*, with violin width scaled to the number of species in each family. c) Posterior distribution of the family-level difference in mean random intercept (*Arenaviridae* − *Hantaviridae*), computed per posterior draw to propagate uncertainty. The credible interval includes zero (posterior mean difference = −0.27, 95% CrI [−0.58, 0.03]), indicating no strong evidence of a systematic difference in baseline detectability between the two viral families.

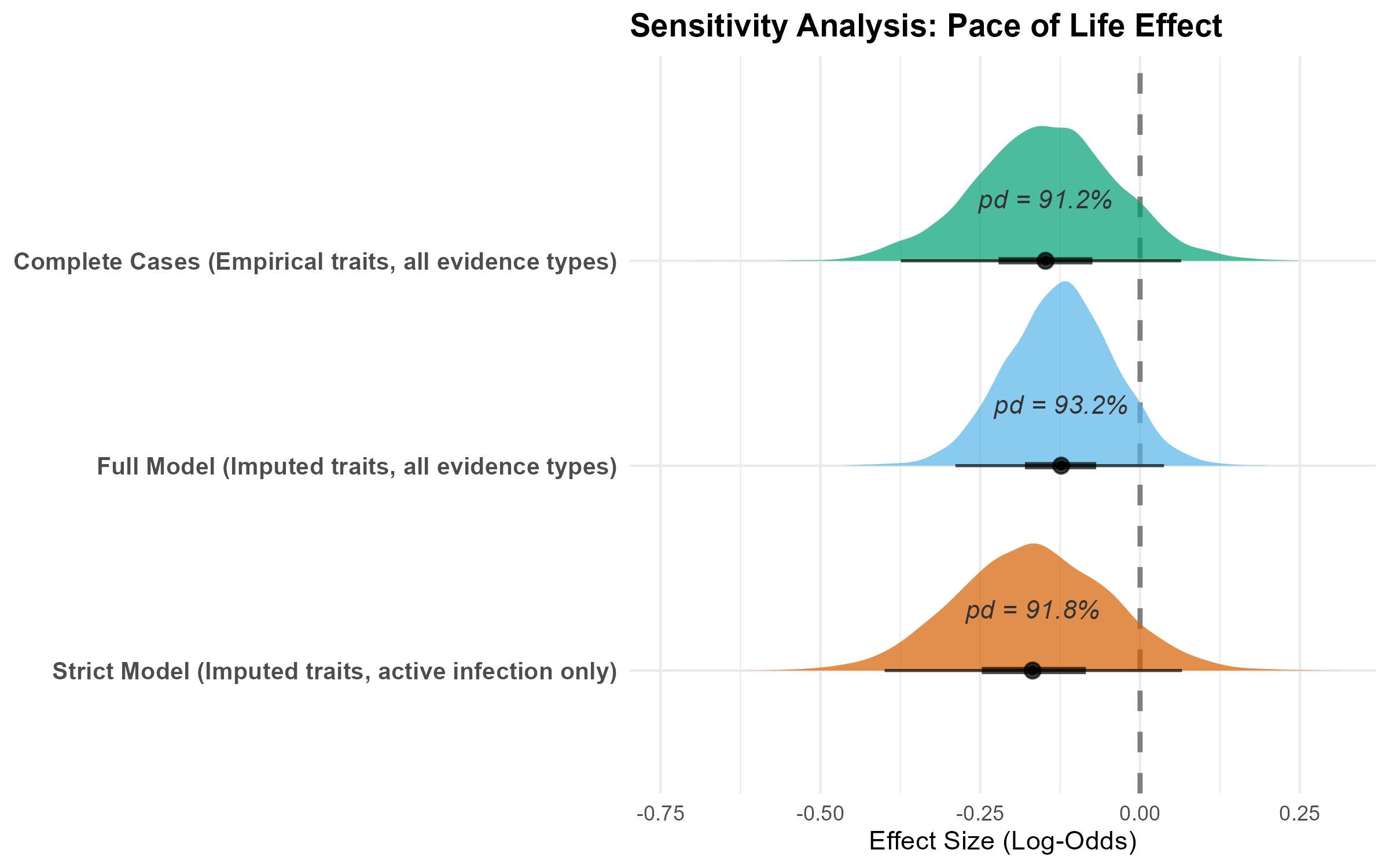

**Supplementary Figure 6 | Sensitivity analysis of the pace-of-life effect.** Posterior log-odds distributions for the Pace of Life (PC1) predictor across three model specifications: complete-case (empirical traits only), full dataset (all evidence types), and strict (active infection only). Directional support is consistent across specifications and remains stable under the strict definition, suggesting the association is not attributable to serological cross-reactivity alone. Credible intervals cross zero in all three models.

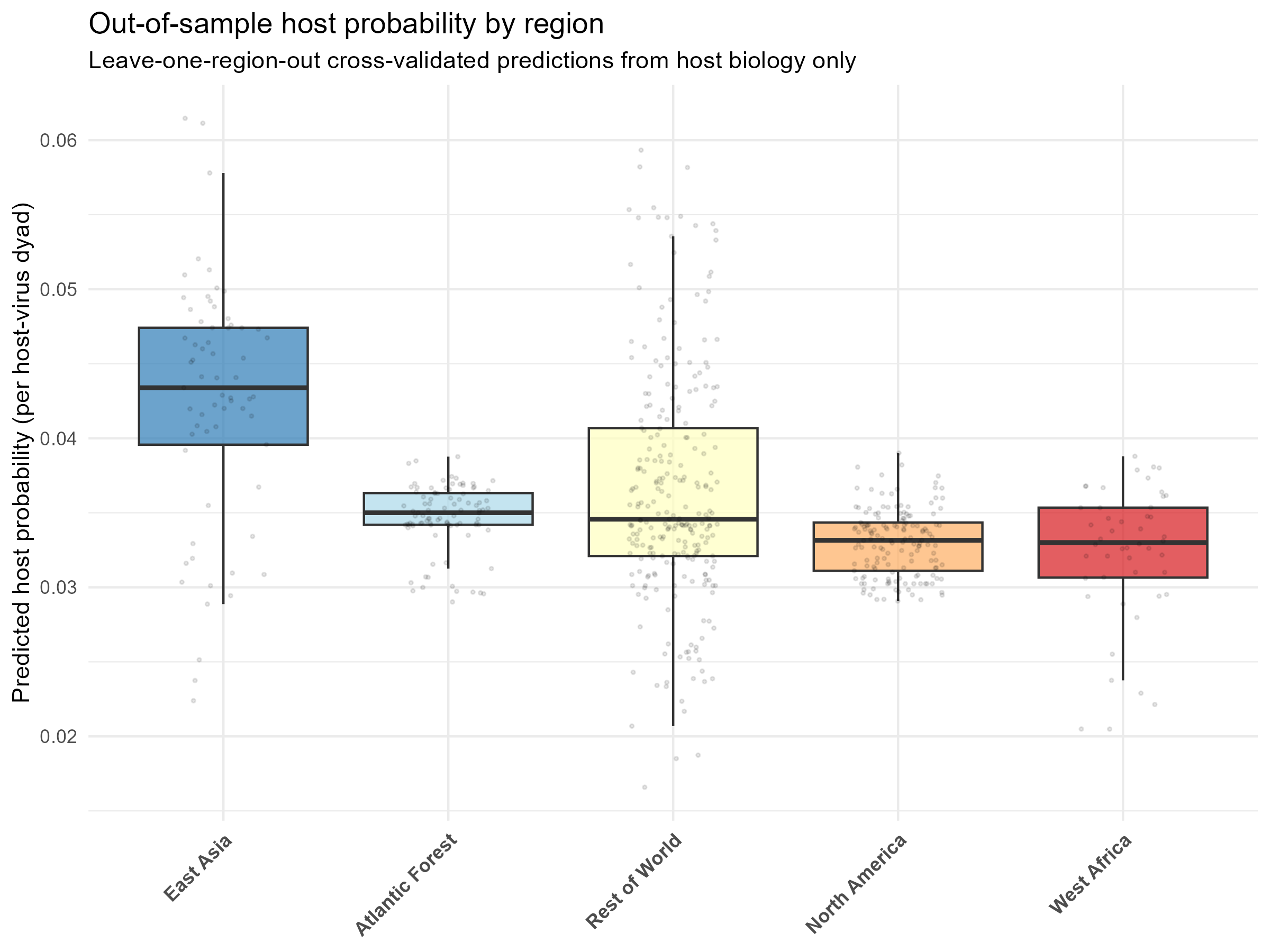

**Supplementary Figure 7 | Regional variation in host probability.** Boxplots displaying the distribution of predicted community host probability for small mammal assemblages across major geographic regions. Predictions are derived from intrinsic pace-of-life traits conducted as part of the cross-fold validation using a leave-one-region-out approach, independent of historical sampling effort or viral identity.

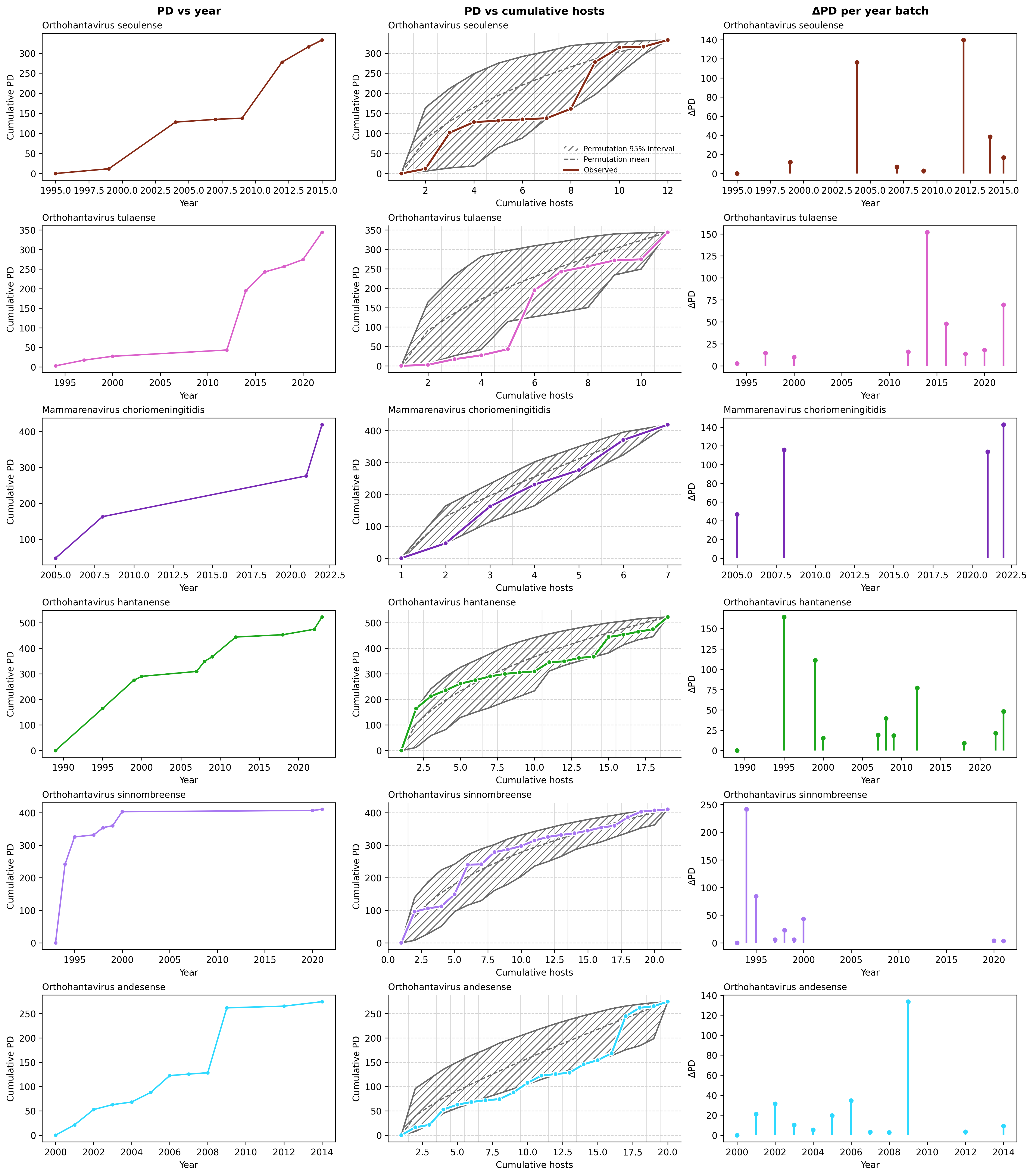

**Supplementary Figure 8 | Accumulation of host phylogenetic diversity for exemplar viruses.** Cumulative host phylogenetic diversity (PD) trajectories for six exemplar viral species (*Orthohantavirus andesense*, *O. hantanense*, *O. seoulense*, *O. sinnombreense*, *O. tulaense* and *Mammarenavirus choriomeningitidis*). Step-like increases mark years in which phylogenetically distinct hosts were first detected.

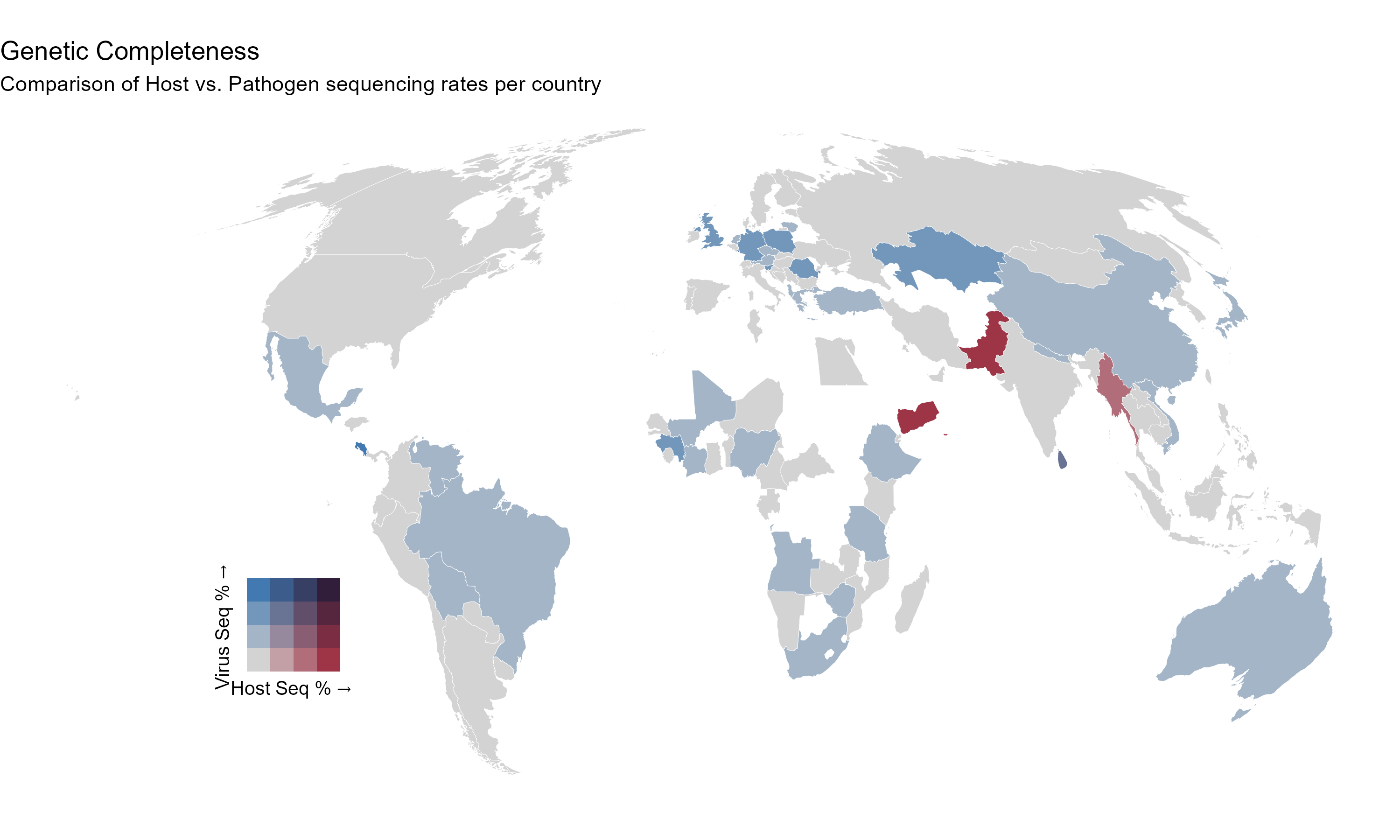

**Supplementary Figure 9 | The geographic genetic completeness gap.** Bivariate map comparing the proportion of sampled hosts that have been sequenced (Host Seq %) versus the proportion of pathogen-positive samples that have yielded valid sequence data (Virus Seq %) aggregated by country.

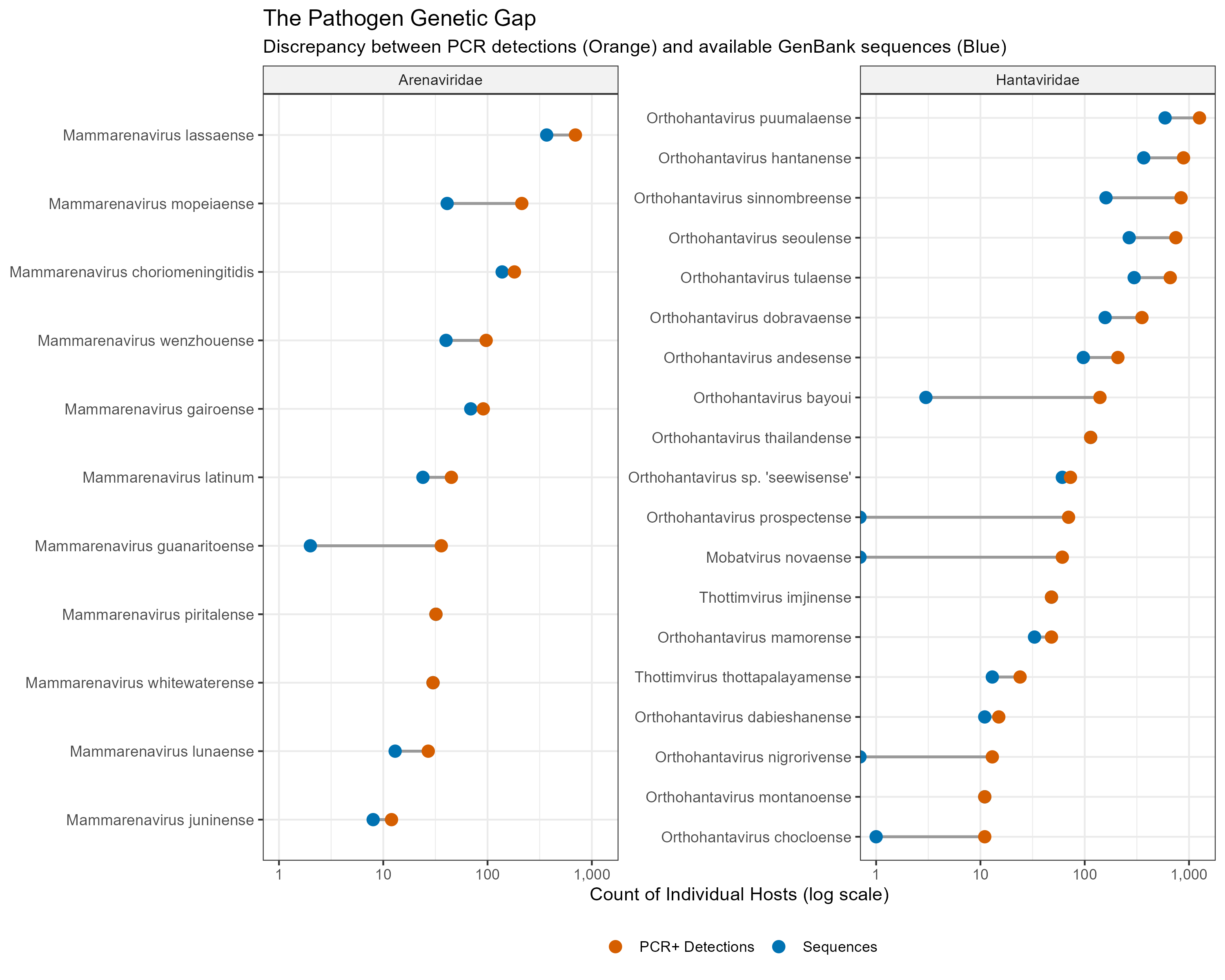

**Supplementary Figure 10 | The pathogen genetic gap.** Dumbbell plot illustrating the discrepancy between the number of individual hosts testing positive via molecular assays (PCR or isolation; orange) and the number of those positive hosts with available GenBank sequences (blue) for each viral species. Viruses are faceted by family.

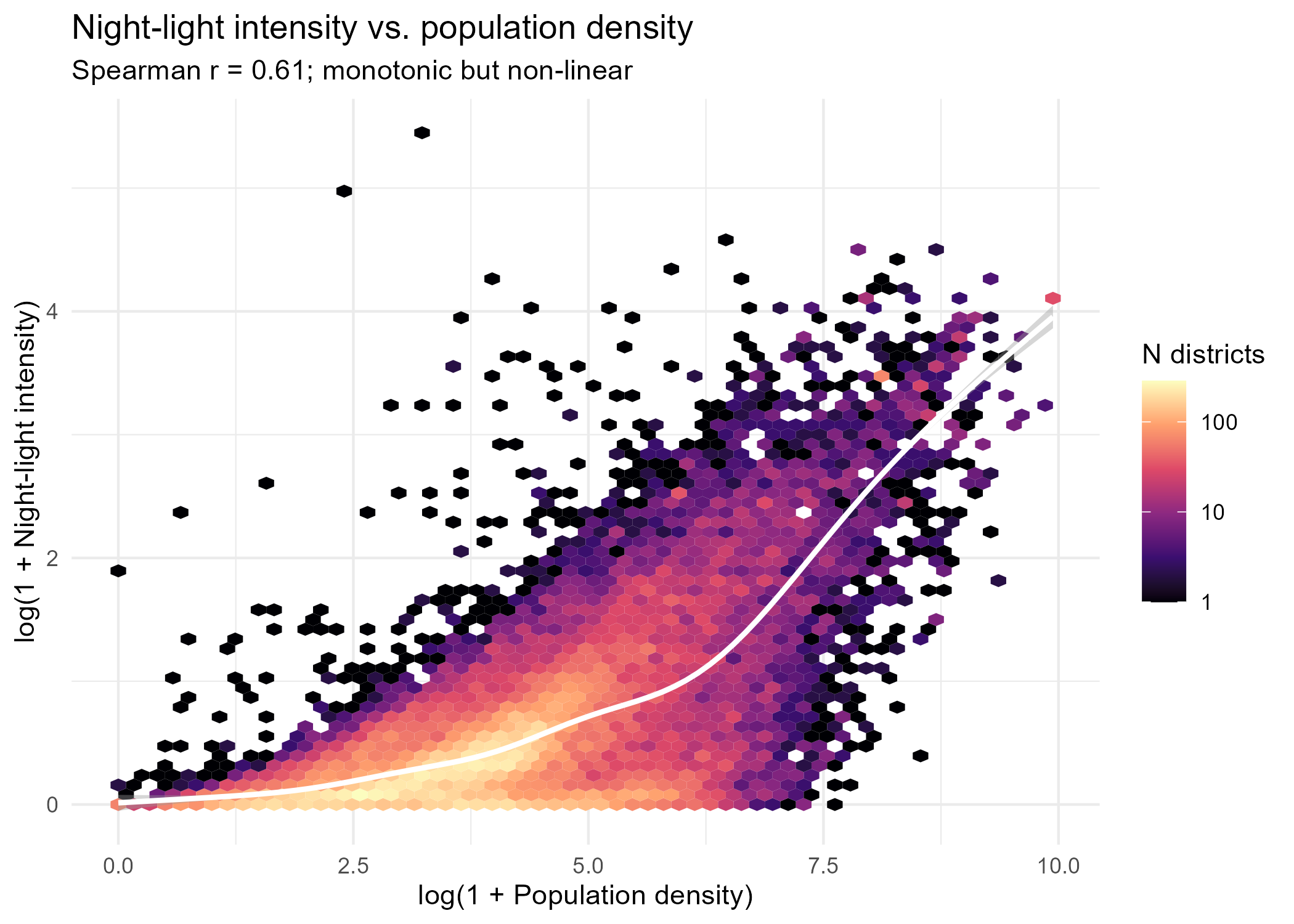

**Supplementary Figure 11 | Relationship between night-time light intensity and population density.** Hexagonal binning of administrative districts (N = 38,897), with a generalised additive model smooth overlaid. The two covariates are positively correlated (Spearman’s ρ = 0.61, p < 0.001) but the relationship is non-linear, with a GAM smooth explaining more deviance than a linear fit (57% vs. 47%). Districts combining high population density with low night-light output are consistent with limited infrastructure relative to population; a small number of districts show the inverse pattern (high light, low density), corresponding to major oil and gas extraction infrastructure rather than settlement.

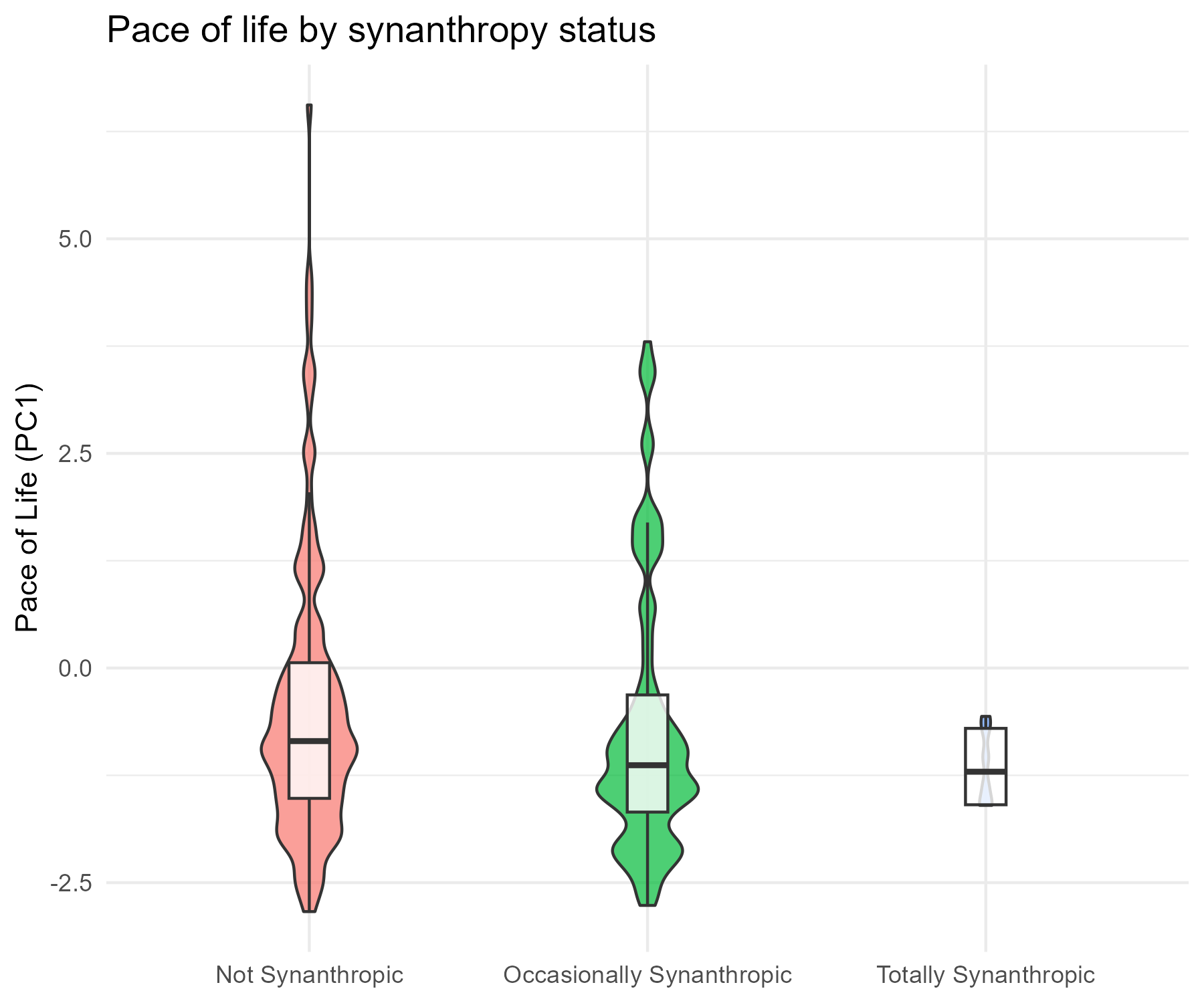

**Supplementary Figure 12 | Pace of life by synanthropy status.** Violin plots showing the distribution of Pace of Life (PC1) scores across synanthropy categories in the subset model (N = 18,906 dyads), with violin width scaled to the number of observations per category and boxplots showing median and interquartile range. Mean PC1 shifts monotonically from Not Synanthropic (−0.53) through Occasionally Synanthropic (−0.76) to Totally Synanthropic (−1.20). The association is detectable (ANOVA F(2, 18903) = 77.4, p < 0.001) but the effect size is small (η² = 0.008), indicating the two traits carry largely non-redundant information.
